# Divergent evolution of the PRPS enzymes across the tree of life

**DOI:** 10.64898/2026.06.01.728777

**Authors:** Bibek R. Karki, Jarek Meller, John T. Cunningham

**Author notes:** Corresponding authors: John T. Cunningham,; Bibek R. Karki.

## Abstract

The phosphoribosyl pyrophosphate synthetase (PRPS) enzyme plays a central role in core biochemical pathways across all life, reflecting its deep evolutionary significance. Here, we present a pan-domain analysis of more than 35,000 non-redundant protein sequences that defines the fundamental features of PRPS at the roots of both domains of life and at critical branchpoints in the tree of life, including during early eukaryogenesis. Combining protein language modeling with maximum likelihood phylogenetic analysis, we identify entirely new PRPS enzyme classes and reveal how neofunctionalization of the canonical class I (bacteria-derived) or class III (archaea-derived) enzymes proceeds via genetic drift or gene duplication. We further demonstrate that multiple PRPS classes from distinct bacterial ancestries were transferred to the eukaryotic genome prior to supergroup radiation, and we provide biochemical and structural characterization of representative examples to clarify their roles in eukaryotic metabolism. Finally, we identify over 30 independent instances of PRPS pseudoenzyme formation across nearly all major eukaryotic lineages and PRPS orthologs, revealing a widespread but underappreciated mechanism of PRPS regulation. Together, this systems-level investigation resolves some of the earliest genetic events shaping life on Earth and offers detailed insight into the evolutionary mechanisms that sculpt enzyme structure and function.

## INTRODUCTION

The phosphoribosyl pyrophosphate synthetase enzyme (PRPS) is the emanation point for multiple biochemical pathways that are universally conserved and predate the last universal common ancestor (LUCA)^1–3^. It utilizes a pyrophosphate from a nucleotide triphosphate to convert ribose-5-phosphate into 5-phosphoribosyl-1-pyrophosphate (PRPP), which is in turn used for the synthesis of purine, pyrimidine, and pyridine nucleotides as well as the amino acids tryptophan and histidine^4,5^. As a “universal enzyme” that also catalyzes one of the rare chokepoint reactions in cellular biochemistry, investigating the ancestry and complete landscape of the PRPS protein universe across the tree of life holds great promise for yielding insights into evolutionary processes and patterns that date from life’s ancient root all the way to the ends of each of its branches^6,7^. However, such reconstruction studies are technically challenging because metabolic subsystem components (unlike those responsible for replicating, transcribing and translating the genetic code, for example) do not tend to follow an easily traceable vertical inheritance pattern due to the gain and loss of paralogs, xenologs, and orthologs.

Here, we deploy orthogonal approaches to disentangle the complex ancestry of PRPS homologs – beginning first with the eukaryotes and then expanding our analysis to Bacteria and Archaea – to determine how the various classes of PRPS enzymes have been generated from a shared ‘urzyme’ ancestral form that existed in a progenitor of the universal common ancestor. We identify eleven ancient forms of the PRPS enzyme whose origins predate radiation of bacteria. We reveal that the eukaryote stem progenitor acquired three of these distinct donated PRPS genes from bacteria, one of which encodes a novel class of PRPS that we characterize for the first time. We pinpoint and order gene duplications in the stem eukaryote progenitor that created the paralogs we observe throughout modern eukaryotes, and we identify numerous additional PRPS gene duplications and losses at key branchpoints during eukaryotic radiation. Lastly, we uncovered a previously unappreciated major role of PRPS pseudoenzymes in regulating PRPS activity across multiple diverse eukaryotic lineages. Together, our work provides an important resource for the field of evolutionary biology that resolves some of the earliest diversification events in the evolution of prokaryotes and eukaryotes, and it provides new examples of how known evolutionary processes can transform an enzyme in both form and function.

## RESULTS

### Acquisition of three distinct ancestral PRPS classes and their expansion and partitioning during eukaryogenesis

To reconstruct the early evolution of PRPS enzymes in eukaryotes, we curated PRPS homologs from representative species spanning every major eukaryotic supergroup (Supplementary Table 1). Because PRPS homologs are deeply conserved yet have undergone repeated duplication, divergence, and loss, we analyzed them with complementary approaches that capture both global sequence similarity and evolutionary relationships: protein language model embeddings, alignment-based pairwise distance analysis, and maximum-likelihood gene tree analysis.

Uniform Manifold Approximation and Projection (UMAP) of Evolutionary Scale Modeling-2 (ESM-2)-derived protein embeddings^8^ resolved eukaryotic PRPS sequences into three discrete clusters: the previously characterized Class I^5,9–15^ and Class II enzymes^16–21^, and a third cluster, clearly separated from both, comprising sequences from organisms in all three major eukaryotic supergroups (Fig.1A). We designate this third cluster as Class IV PRPS Class III has already been assigned to archaeal PRPS^5,22–24^. An orthogonal alignment-based approach, in which sequences were projected by UMAP from a pairwise distance matrix, recovered the same three clusters and further resolved Class I into four subclasses and Class II into two subclasses (Fig.1B). These separations were further supported by silhouette analysis of unsupervised spectral clustering (Supplementary Fig.1A-C) as well as an inferred maximum-likelihood gene tree (Fig.1C, Supplementary Figs.1D-E, and Supplementary File 1).

**Figure 1.**
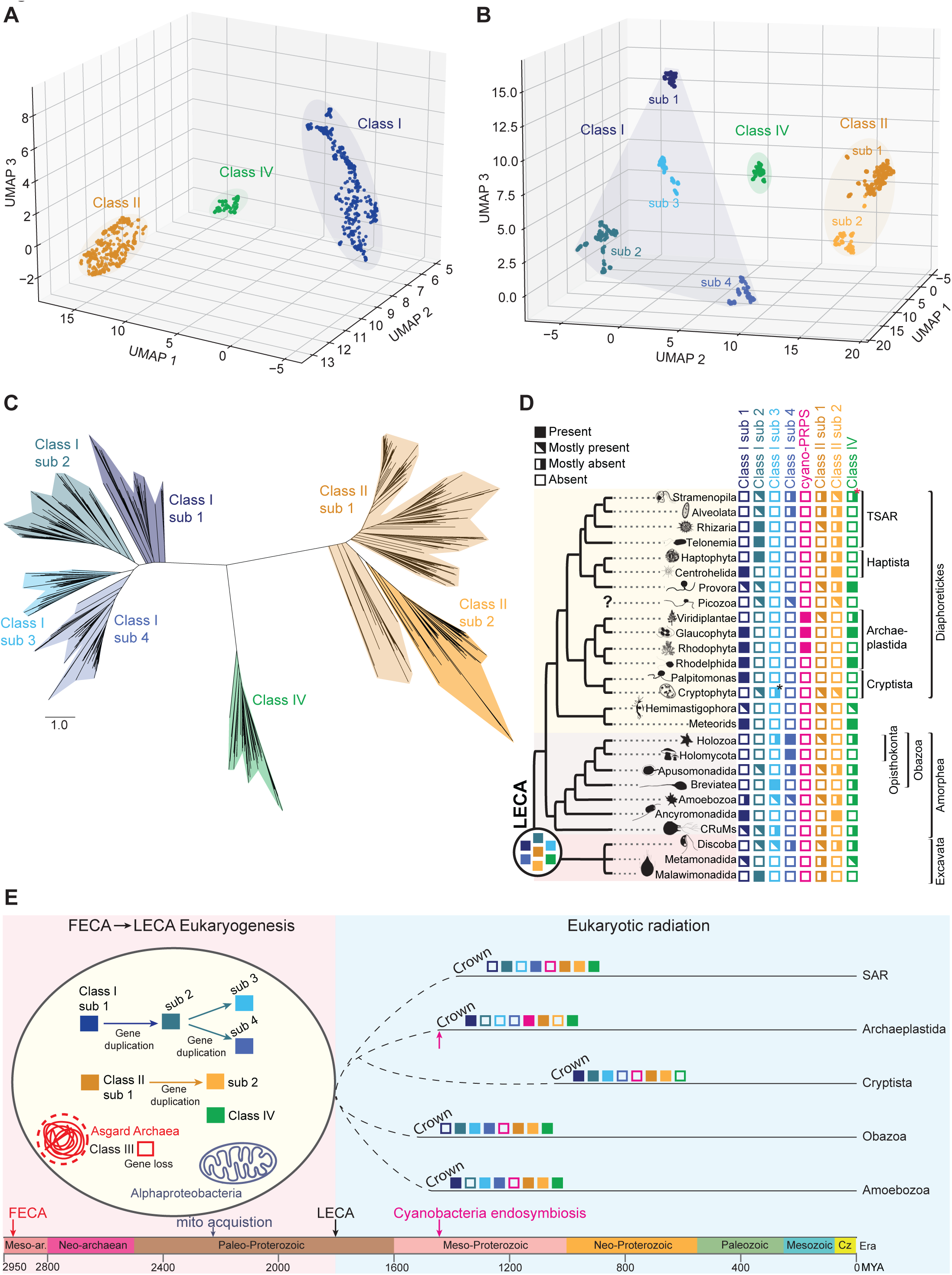
Acquisition of three distinct ancestral PRPS classes and their expansion and partitioning during eukaryogenesis. **(A)** Three-dimensional UMAP of ESM-2-derived protein embeddings for eukaryotic PRPS sequences (n = 802) from representative species across all eukaryotic supergroups, colored by class assignment and projected using cosine distance. **(B)** Three-dimensional UMAP of eukaryotic PRPS sequences based on a pairwise p-distance matrix derived from a MAFFT-generated multiple sequence alignment. **(C)** Maximum likelihood gene tree of eukaryotic PRPS sequences inferred using IQ-TREE under the LG+F+R10 model. The full tree is provided in rectangular format with bootstrap support values in Supplementary File 1 and in Newick format in the Figshare repository. **(D)** Distribution of PRPS enzyme classes and subclasses across major eukaryotic lineages. “Mostly present” indicates that a given class is detected in the majority of sampled taxa within a lineage, whereas “mostly absent” indicates limited representation. “Absent” denotes no detectable homologs in the available datasets. A detailed lineage-level breakdown is provided in Supplementary Figure 5. “cyano-PRPS” corresponds to cyanobacteria-derived PRPS acquired via primary endosymbiosis in the common ancestor of Archaeplastida. Black asterisk denotes Class I subclass 3 PRPS restricted to a single Cryptophyta species (uncultured Katablepharidaceae), consistent with horizontal gene transfer, and red asterisk indicates Class IV PRPS in a limited number of Stramenopila (three Xanthophyceae species), also suggestive of horizontal gene transfer. Question mark next to Picozoa denotes uncertain placement within Diaphoretickes. **(E)** Model for PRPS class evolution during eukaryogenesis. Three ancestral PRPS genes (Class I, Class II, and Class IV) were present in the stem eukaryote, with subsequent duplications during the FECA-LECA transition expanding Class I into four subclasses and Class II into two, while Class IV remained a distinct single-copy gene. Following LECA, eukaryotic radiation was accompanied by extensive, lineage-specific gene loss, resulting in differential retention of PRPS classes across extant lineages. An additional PRPS gene (cyano-PRPS) was acquired in Archaeplastida via primary endosymbiosis, while archaeal Class III PRPS was lost during early eukaryogenesis. Estimated evolutionary timeline adapted from references^6,27^.

To determine whether these class and subclass designations reflected position-specific sequence signatures rather than global similarity alone, we identified diagnostic residues that were enriched within, and depleted outside, each class or subclass. Each class and subclass possessed a distinct diagnostic residue panel (Supplementary Figs.2A-C). Several diagnostic residues mapped to known functionally important regions of PRPS enzymes, whereas others occurred at residues of currently undetermined significance (Supplementary Fig.3). To ascertain whether these residues nevertheless imbue functional significance, we examined PRPS1 and PRPS2 – a Class I paralog generated by duplication of PRPS1 in the Gnathostomata stem^25^. The diagnostic residues distinguishing PRPS1 and PRPS2 fell outside canonical functional regions (Supplementary Figs.4A and 4B), but nonetheless impart functional differences in their biochemistry and respective abilities to sculpt assembly and activity of the heteromeric mammalian PRPS complex^14,25,26^. Thus, the diagnostic residues of currently undetermined significance across eukaryotic PRPS classes and subclasses may similarly encode functional differences

The distribution of PRPS across the eukaryotic phylogenetic tree^27–29^ showed that Class I PRPS was nearly universally retained, whereas Class II and Class IV displayed sporadic and broadly mutually exclusive distributions, with co-occurrence confined to a few Archaeplastida and Provora species (Fig.1D and Supplementary Fig.5). Despite this sporadic distribution, both Class II and Class IV were detected across eukaryotic supergroups, a pattern consistent with their presence in an early eukaryotic ancestor followed by repeated lineage-specific losses. Within Archaeplastida, we also identified an additional Class I-like subset that was restricted to glaucophytes, red algae, and green plants, and possessed significant similarity cyanobacterial Class I sequences. This combination of phylogenetic distribution and sequence similarity is most parsimoniously explained by primary endosymbiotic gene transfer as has previously been suggested^30^. We refer to these Archaeplastida-restricted sequences as cyano-PRPS hereafter, to distinguish them from other Class I sequences whose origins predate the Last Eukaryotic Common Ancestor (LECA).

Class I and Class II subclass members were detected across multiple eukaryotic supergroups, indicating that these PRPS paralogs were already established before the radiation of extant eukaryotes from LECA. To distinguish ancient paralogs from independently acquired genes, we examined subclass co-existence within individual genomes. Several species from distantly related eukaryotic lineages retained multiple Class I or Class II subclasses, and these sequences grouped with their respective subclass clades in the eukaryotic PRPS gene tree rather than arranging by established phylogenetic relationships (Supplementary Figs.1D and 1E). This pattern indicates that, for both Class I and Class II, the subclasses represent ancient paralogs rather than products of recent lineage-specific expansion.

To further confirm the shared ancestry of these PRPS paralogs, we compared exon-intron architectures across representative orthologs, taking advantage of splice-junction conservation over deep evolutionary timescales^31,32^. Multiple splice junctions were conserved across Class I subclasses and across Class II subclasses that spanned eukaryotic supergroups which was a pattern also observed in Class IV sequences (Supplementary Figs.6 and 7). These distinct conserved junctions support common ancestry among subclasses within each class while confirming Class I, Class II, and Class IV were separate ancestral PRPS genes. Within Class I, the strongest splice-junction sharing occurred between subclasses 1 and 2, with subclass 2 also sharing junctions with subclasses 3 and 4 (Supplementary Fig.6B). This nested pattern is consistent with subclass 1 representing the earliest-diverging Class I subclass, followed by subclass 2, with subclasses 3 and 4 arising through subsequent duplications from a subclass 2-like ancestor. Taken together, the converging evidence – that sequences sampled across all three major eukaryotic supergroups fall into the same Class I and Class II subclasses, that multiple subclasses co-exist within single genomes of distantly related eukaryotes, and that conserved splice-junction patterns are shared both within and between subclasses – supports pre-LECA duplication of Class I into four paralogs and Class II into two paralogs, alongside a single-copy Class IV-encoding gene.

To nominate candidate bacterial sources and further resolve the relative order of subclass emergence, we reconstructed ancestral sequences for LECA Class I subclasses 1-4, LECA Class II subclasses 1-2, LECA Class IV, and Ancestral Cyanobacteria-derived PRPS (ACD-PRPS), and queried each by BLASTP against 33,248 non-redundant prokaryotic PRPS sequences (Supplementary Tables 2 and 3). The resulting hit profiles differed among reconstructed ancestors (Supplementary Fig.8). As expected, nearly all top BLASTP matches for ACD-PRPS were cyanobacterial, consistent with its endosymbiotic origin^33–35^. Among Class I reconstructions, subclass 1 showed the strongest similarity to prokaryotic PRPS sequences, with higher top-hit bit scores than subclasses 2-4; similarly, Class II subclass 1 was more closely matched to prokaryotic sequences than subclass 2. These results independently corroborate respective subclass 1 genes as the least divergent orthologs in both Class I and Class II^36^, nominating them as the original genes acquired from bacteria during eukaryogenesis. LECA Class I, Class II, and Class IV each showed strongest BLASTP similarity to distinct bacterial lineages, while archaeal and alphaproteobacterial sequences were not among their top-percentile matches. Thus, the pre-LECA eukaryotic PRPS repertoire was not inherited vertically from the archaeal host lineage, nor was it acquired via horizontal gene transfer (HGT) from the alphaproteobacterial-derived mitochondrial endosymbiont^7,37–43^.

Together, these observations support the model summarized in Fig. 1E^6,29^. Three distinct PRPS genes – the ancestors of eukaryotic Class I, Class II, and Class IV – were acquired from non-alphaproteobacterial sources before LECA. During the FECA-to-LECA transition, Class I expanded from a subclass 1 ancestor through stepwise duplications that first generated subclass 2 and subsequently gave rise to subclasses 3 and 4, whereas Class II expanded from a subclass 1 ancestor to generate subclass 2. Together with a single-copy Class IV gene, these events established a seven-gene PRPS repertoire in LECA. A subsequent, lineage-restricted acquisition of cyano-PRPS through primary endosymbiosis added an additional PRPS gene to Archaeplastida. Following LECA radiation, this expanded repertoire was differentially partitioned across descendant taxa through extensive lineage-specific gene loss, which stripped away much of LECA’s metabolic redundancy. This process produced the heterogeneous and broadly mutually exclusive distribution of Class II and Class IV, alongside the near-universal retention of Class I, observed in extant eukaryotes.

### Expansion of PRPS classes during bacteriogenesis supports a metabolically rich ancestral bacterial pangenome

Our previous analyses suggested that the three original pre-LECA eukaryotic PRPS genes encoding different classes were acquired from bacterial sources distinct from both the archaeal host and the alphaproteobacterial-derived mitochondrial endosymbiont. However, the broader prokaryotic PRPS landscape, and the bacterial lineages most likely to have contributed to the early eukaryotic repertoire, remained unresolved. Bacterial PRPS has historically been viewed as a near-universal Class I enzyme^44–48^, whereas archaeal PRPS has been represented by a distinct Class III enzyme^5,22–24,49^. To place the inferred eukaryotic ancestors within this broader context, we extended the same comparative workflow used for eukaryotic PRPS to bacterial and archaeal sequences sampled across diverse prokaryotic taxa (Supplementary Tables 2 and 3).

UMAP of ESM-2-derived embeddings revealed a substantially more diverse prokaryotic PRPS landscape than previously known (Fig.2A). Bacterial Class I and archaeal Class III sequences formed dense, dominant clusters, but multiple discrete clusters fell outside these canonical classes, including bacterial Class II and Class IV, which were identified by sequence similarity and diagnostic residues shared with their eukaryotic counterparts. We used this landscape as a discovery framework to define the major clusters as distinct PRPS classes, expanding the prokaryotic classification beyond Class I to include bacterial Classes II and IV-XI, and beyond Class III to include an additional archaeal Class XII. Class I and Class III PRPS were distributed across all major bacterial and archaeal lineages, respectively, whereas other classes were distributed more sporadically (Supplementary Figs.9A and 9B). The landscape also revealed a region comprised of heterogenous sequences with cross-domain overlap. Interestingly, the archaeal sequences in this region as well as those found within the bacterial Class I cluster were predominantly from the Nanobdellati lineage (formerly DPANN archaea), and most of the bacterial sequences in the heterogenous region were predominantly from Candidate Phyla Radiation (CPR). Both Nanobdellati and CPR are comprised of microorganisms characterized by ultrasmall cells, reduced genomes, symbiotic lifestyles, limited biosynthetic capacity, variable metabolic repertoires and a history of acquiring proteins via ancient HGT from diverse prokaryotes^50–57^. Overlaying ancestrally reconstructed eukaryotic sequences onto this space placed ACD-PRPS and LECA Class I subclass 1 within bacterial Class I, whereas LECA Class II subclass 1 and LECA Class IV localized within bacterial Class II and Class IV clusters, respectively (Fig.2A).

**Figure 2.**
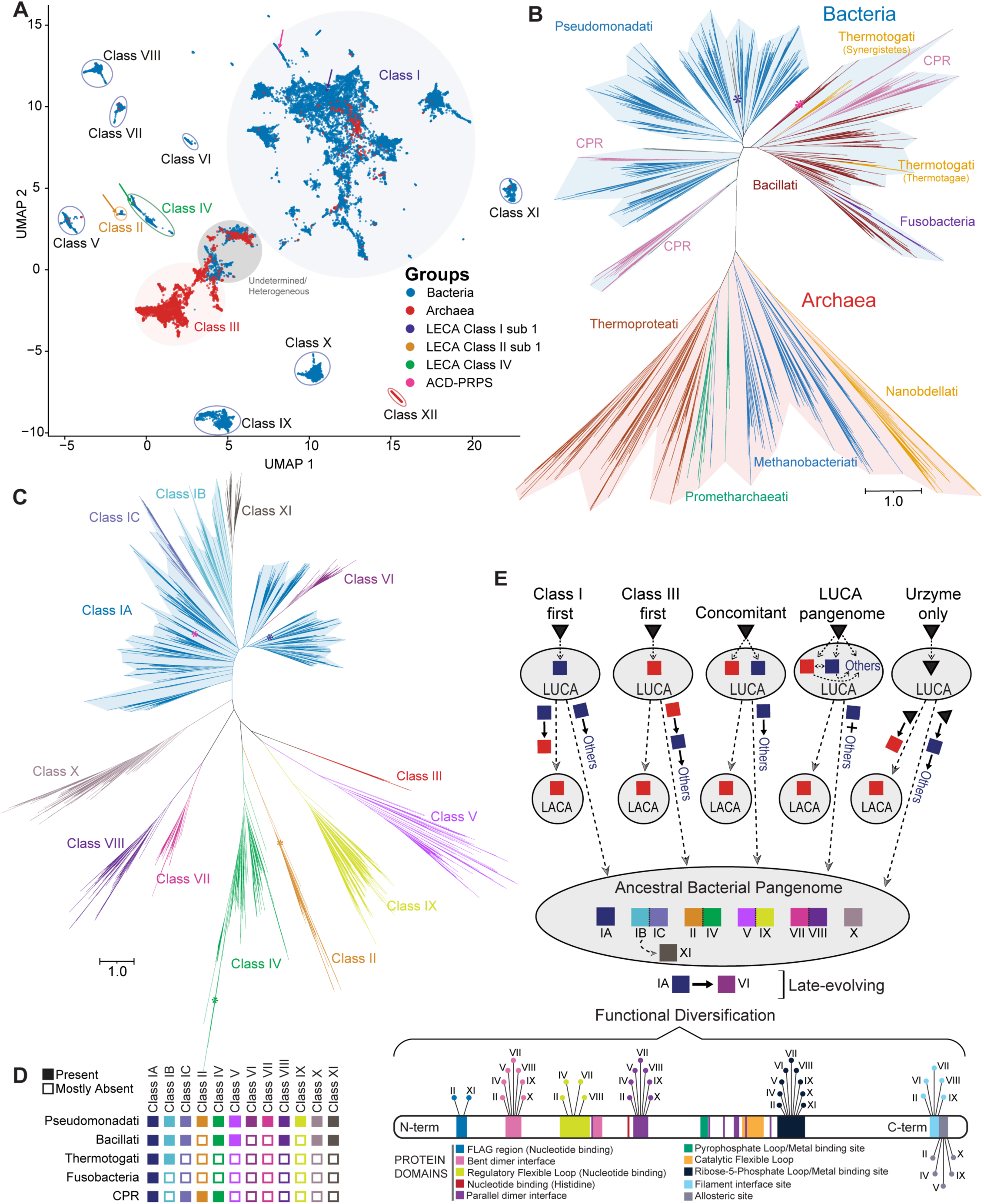
Expansion of PRPS classes during bacteriogenesis supports a metabolically rich ancestral bacterial pangenome. **(A)** Two-dimensional UMAP of ESM-2-derived embeddings for bacterial (n = 29,322) and archaeal (n = 4,686) PRPS sequences, colored by domains. Each point represents a single sequence. Clusters outside canonical bacterial Class I and archaeal Class III are circled and designated as additional PRPS classes. Colored arrows denote ancestrally reconstructed eukaryotic sequences (blue, LECA Class I sub 1; pink, ACD-PRPS; orange, LECA Class II sub 1; green, LECA Class IV). **(B-C)** Maximum likelihood gene trees of PRPS sequences inferred using IQ-TREE under the LG+F+R10 model. (B) Gene tree of bacterial Class I and archaeal Class III sequences, with branches colored by major prokaryotic lineages. (C) Expanded tree including additional bacterial and archaeal PRPS classes, with branches colored by PRPS class. The full tree is provided in rectangular format with bootstrap support values in Supplementary Files 4 and 5, respectively, and in Newick format on the Figshare repository. Asterisks mark positions of ancestrally reconstructed eukaryotic sequences. **(D)** Distribution of bacterial PRPS classes across major kingdoms and CPR. **(E)** Evolutionary models for PRPS emergence and expansion in prokaryotes. Alternative scenarios for LUCA composition are shown, differing in whether it encoded a Class I-like enzyme (Class I first), a Class III-like enzyme (Class III first), both Class I and Class III (Concomitant), a more complex PRPS repertoire comprising multiple classes (LUCA pangenome), or a minimal urzyme (Urzyme only). Across all scenarios, the Last Archaea Common Ancestor (LACA) inherits a Class III-like PRPS enzyme, whereas the bacterial stem expands early to generate a diverse, multi-class PRPS repertoire, which is subsequently partitioned across bacterial lineages. Together, these models converge on the presence of a metabolically rich ancestral bacterial pangenome comprising multiple PRPS classes (IA-II, IV, V, and VII-XI), with an additional Class VI arising through later lineage-specific innovation from Class IA. The lower panel summarizes functional diversification by mapping changes across key structural and regulatory features relative to Class I, including nucleotide-binding regions, flexible loops, metal-and substrate-binding sites, and oligomerization interfaces.

**Figure 3.**
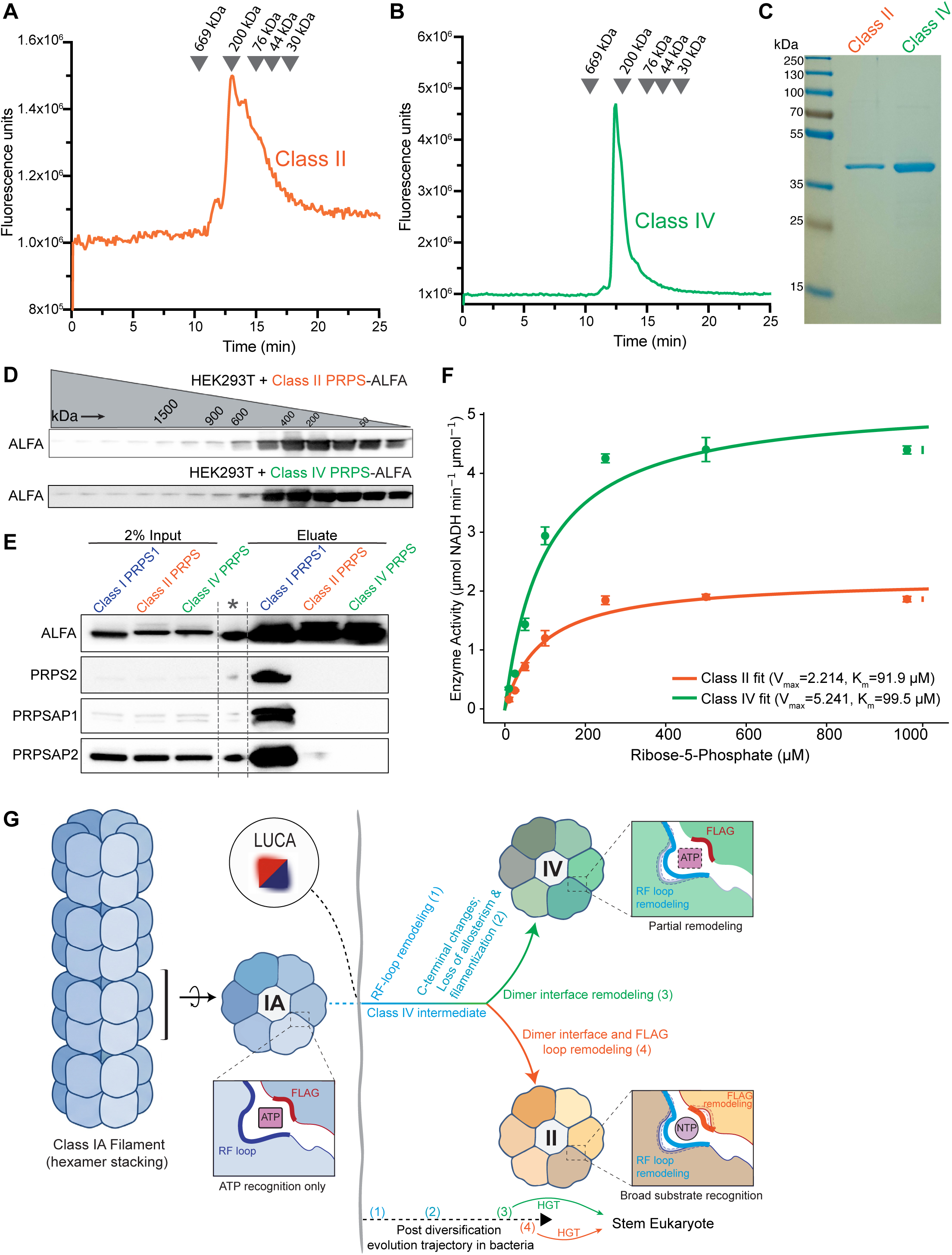
Functional and structural divergence of Class II and Class IV PRPS enzymes supports a stepwise evolutionary trajectory from a Class I-like ancestor. **(A-B)** Size-exclusion chromatography (SEC) elution profiles of bacterially purified recombinant Class II PRPS from *Branchiostoma lanceolatum* (A) and Class IV PRPS from *Corallochytrium limacisporum* (B) on a Yarra SEC-2000 column. Reference molecular weight standards are shown. **(C)** SDS-PAGE with Coomassie staining of peak SEC fractions from (A-B). **(D)** Western blot analysis of SEC fractions from HEK293T cells heterologously expressing ALFA-tagged Class II (*B. lanceolatum)* or Class IV (*C. limacisporum)* PRPS **(E)** SDS-PAGE analysis of eluates from ALFA-tag immunoprecipitation from HEK293T cells expressing C-terminally tagged Class I (*H. sapiens*), Class II (*B. lanceolatum)* and Class IV (*C. limacisporum)* PRPS. Asterisk indicates empty lane with residual signal from adjacent lane spillover. **(F)** Enzyme activity of recombinant Class II and Class IV PRPS from peak fractions shown in (A-B) across increasing ribose-5-phosphate concentrations, measured via PRPS activity assay (Supplementary Figure 19). Activity is reported as µmol NADH min⁻¹ µmol⁻¹ enzyme. Data represent mean ± SD (n = 3), with nonlinear least-squares fits to the Michaelis–Menten equation used to derive kinetic parameters (V_max_ and K_m_). **(G)** Schematic of a proposed stepwise evolutionary trajectory linking Class IA, Class IV, and Class II PRPS enzymes. Class IA forms filaments via hexamer stacking; zoomed inset of a representative hexamer highlights dimeric pocket, where ATP recognition involves the RF loop and FLAG region. In this framework, early Class IV represents an intermediate state that retains partial Class I-like features while undergoing (1) RF-loop remodeling and (2) changes at the C-terminus associated with loss of allosteric regulation and filament formation. Subsequent steps include (3) divergence of bent dimer interface residues (Class IV) and (4) additional remodeling of dimer interface and FLAG-region residues (Class II), yielding Class II generalist enzymes with broader substrate recognition. Numbers indicate inferred stepwise transitions. The inclusion of LUCA denotes an alternative scenario in which PRPS diversification may originate from a more ancestral state instead of Class IA to acknowledge that the precise evolutionary ancestor of these classes remains unresolved. Following bacterial diversification, Class II and Class IV PRPS were acquired by the stem eukaryote through horizontal gene transfer (HGT), and their defining sequence features are retained in respective extant eukaryotic sequences.

To probe the ancestral relationships of these classes, we performed a prokaryotic PRPS gene tree analysis of representative bacterial Class I and archaeal Class III PRPS sequences. The inferred maximum-likelihood gene tree separated bacterial and archaeal PRPS enzymes and showed that the most prevalent bacterial Class I sequences mostly followed a vertical inheritance pattern where the sequence groupings in the gene tree largely reflected known taxonomic hierarchies^58^ (Fig.2B, Supplementary Fig.10A and Supplementary Files 2 and 3). We refer to this most evolutionarily successful bacterial Class I gene as Class IA. Of note, PRPS sequences from CPR did not group together to form a single clade within this gene tree, which may reflect differential retention, gene exchange, or the limited resolution of a single-gene phylogenetic construction^59^ in addition to their ancient, diverse and symbiotic nature^52–57^. Thus, the origin and inheritance of Class IA PRPS within CPR remain unresolved. Similar to Class IA, archaeal Class III sequence clustering in the gene tree largely reflected known taxonomic hierarchies in that domain^58^, whereas Class XII formed a distinct clade predominantly consisting of sequences from extreme halophiles^60^ within Methanobacteriota (Fig.2B, Supplementary Fig.10B and Supplementary Files 2 and 4). Collectively, these results establish Class IA and Class III as the most prevalent PRPS enzymes – each following a clear vertical gene inheritance pattern in Bacteria and Archaea, respectively.

We curated Class IA on the basis of its wide distribution and vertical inheritance pattern, but a subset of sequences within the Class I embedding region did not follow the expected bacterial taxonomic hierarchies^58^ when included in the preliminary PRPS Class I gene trees. Two such conserved groups, designated Class IB and Class IC, were detected across multiple bacterial kingdoms, supporting ancient diversification within Class I. To resolve these Class I subclasses (Class IA, Class IB, and Class IC) alongside the additional PRPS classes identified by ESM-2 protein language modeling, we inferred an expanded maximum-likelihood gene tree (Fig.2C, Supplementary Fig.11 and Supplementary File 5). This tree separated bacterial PRPS into Class I – comprising Class IA, IB, and IC – and additional Classes II and IV-XI. Class IB and Class IC formed distinct subclades within bacterial Class I and occupied separate positions within the broader Class I landscape (Supplementary Fig.12). Several pairs of classes – Class II and Class IV, Class V and Class IX, Class VII and Class VIII, and Class IB and Class IC – formed sister relationships in the gene tree, consistent with ancient divergence from common ancestors. Class XI was nested within the Class IB group despite falling outside the broader Class I cluster in embedding landscape, indicating that a subclass IB ancestor gave rise to a distinct PRPS class. Likewise, Class VI appeared as a later taxon-restricted expansion directly from Class IA within Rhodobacterales (Alphaproteobacteria). When ancestrally reconstructed eukaryotic sequences were placed onto this expanded PRPS gene tree, each localized near a distinct bacterial group rather than within Alphaproteobacteria or the archaeal Class III clade (Fig.2C and Supplementary Fig.11).

Distribution of these classes across Bacteria revealed a disparate pattern of inheritance and retention (Fig. 2D). Class IA was broadly distributed, whereas most other PRPS classes were more restricted, with some spanning multiple bacterial kingdoms and others confined to a single kingdom. This pattern is consistent with differential retention and loss from a broader ancestral bacterial PRPS repertoire rather than repeated independent origin of each class in disparate bacterial lineages. However, we cannot rule out the possibility that ancient HGT may have contributed to the sporadic distribution of some classes, which confounds strict interpretation of vertical inheritance. Bacterial Class II and Class IV were mutually exclusive within genomes, mirroring their complementary distribution in eukaryotes. However, these classes often co-existed in genomes alongside Class I rather than replacing it – 52.7% of Class II-harboring species and 74% of Class IV harboring species also possessed a Class I PRPS gene (Supplementary Table 2) – nominating Class II and Class IV as professional augmenters of Class I.

To determine whether the prokaryotic PRPS classes differed at functionally relevant positions, we examined conservation and diagnostic residue patterns. Full-length sequence logos showed that universally conserved catalytic residues were retained across major PRPS classes, consistent with strict preservation of the urzyme-derived PRPP synthetase reaction. (Supplementary Fig. 13). Distinct but related amino acid signatures were also observed between the sister classes identified in gene tree analysis, further supporting their shared ancestry. Importantly, class-specific conserved differences were observed across regions involved in nucleotide binding, substrate coordination, allosteric regulation, and oligomerization interfaces. Diagnostic-residue analysis further confirmed that prokaryotic PRPS classes and bacterial Class I subclasses possess distinct residue signatures (Supplementary Figs. 14A and 14B). This indicates that the prokaryotic classes defined by embedding landscape and gene tree analysis also harbor class-specific sequence features that reflect divergence in structural, regulatory, and/or substrate-recognition properties.

To refine candidate donor assignments for the three pre-LECA eukaryotic classes, we performed profile Hidden Markov Model (HMM)^61^ searches of each ancestrally reconstructed sequence against bacterial and archaeal PRPS profiles (Supplementary Figs. 15A-D). For ACD-PRPS, the top-scoring profile was Cyanobacteria, consistent with its endosymbiotic origin, and thereby confirming the utility of this approach. LECA Class I subclass 1 scored highest against the Methylomirabilota and Rokubacteria profiles, with the alphaproteobacterial profile ranking substantially lower, further corroborating a non-protomitochondrial origin for the eukaryotic Class I ancestor. LECA Class II subclass 1 and LECA Class IV scored highest against Thermodesulfobacteria and CPR profiles, respectively. Across all four ancestral sequences, archaeal profiles ranked outside the top-scoring matches, providing another line of evidence against vertical inheritance of the pre-LECA eukaryotic PRPS genes from the archaeal host.

Together, these observations support the model summarized in Fig. 2E. In this model, the bacterial stem progenote or ancestral pangenome possessed a metabolically rich assortment of PRPS genes encoding multiple PRPS classes, including Classes IA-II, IV, V and VII-XI, which were subsequently partitioned across descendant bacterial lineages through differential retention and loss, mirroring the phenomenon observed in eukaryotes. Class VI arose later through a lineage-restricted innovation from Class IA within Rhodobacterales, potentially associated with occupying phytoplankton-derived carbon niches^62–64^. This model is compatible with several alternative scenarios for the LUCA-to-stem bacteria transition (Fig.2E, top), and accounts for the central observation that distinct bacterial PRPS sources – inherited from an ancestral bacterial pangenome – contributed to each of the three pre-LECA eukaryotic PRPS classes. By contrast, archaeal PRPS diversity was dominated by Class III, with Class XII representing a halophile-restricted archaeal innovation. Thus, while archaeal PRPS diversity remained limited, bacteria inherited and/or generated a broad reservoir of PRPS genes from which the three ancestral eukaryotic PRPS genes were acquired and expanded.

### Functional and structural divergence of Class II and Class IV PRPS enzymes supports a stepwise evolutionary trajectory from a Class I-like ancestor

Building on our identification of three pre-LECA eukaryotic PRPS classes – with Class II and Class IV forming a sister pair in our gene tree analysis (Fig.2C) – we next examined how these two classes, particularly the novel Class IV, relate biochemically and structurally to well-characterized Class I. To evaluate whether these classes encode functionally active enzymes and to test the structural relationships predicted by their sequence divergence from Class I, we first cloned representative Class II and Class IV proteins from the nearest mammalian relatives encoding each class – *Branchiostoma lanceolatum* (a cephalochordate; Class II) and *Corallochytrium limacisporum* (a pluriformea; Class IV).

Bacterially expressed and purified recombinant Class II and Class IV PRPS eluted predominantly in the dimer-to-hexamer range on Superose 6 size exlusion chromatography (SEC) column (5 kDa to 5 MDa fractionation range; Supplementary Figs.16A-D). To better resolve smaller oligomeric states, the major Superose 6 peaks were re-injected onto a Yarra SEC-2000 column (1 kDa to 300 kDa fractionation range), where Class IV PRPS eluted as a symmetrical peak within the hexameric range, whereas Class II PRPS eluted as an asymmetrical peak consistent with a mixture of oligomers ranging from dimers to hexamers as shown in previous studies from plant Class II PRPS^18–21^ (Figs.3A-C). To test whether this organization is maintained in eukaryotic cells, heterologous ALFA-tagged Class II and Class IV PRPS were overexpressed in mammalian cells and analyzed by analytical SEC. Both proteins were detected predominantly in fractions consistent with dimeric to hexameric assemblies (Fig.3D), indicating that hexameric organization, but not higher-order filament formation, is also maintained in cells. The absence of filament formation in Class II and Class IV PRPS is consistent with sequence-level divergence at the C-terminal hexamer-stacking interface that has been implicated in Class I PRPS filament assembly in prokaryotes^65^ and eukaryotes^66^. Eukaryotic Class I PRPS sequences strongly conserve the C-terminal residues that form this filament interface, whereas the corresponding residues in Class II and Class IV are not conserved or are absent (Supplementary Fig.16E).

We then tested whether these enzymes could co-assemble with the endogenous human Class I PRPS complex. ALFA-tag immunoprecipitation of C-terminally tagged human PRPS1 recovered endogenous PRPS2 and the associated proteins PRPSAP1 and PRPSAP2, consistent with the established mammalian Class I PRPS complex^25^. In contrast, ALFA-tagged Class II and Class IV PRPS did not co-associate with endogenous PRPS1, PRPS2, PRPSAP1, or PRPSAP2 (Fig.3E), indicating that Class II and Class IV have diverged to form class-specific assemblies that do not permit interaction with Class I PRPS. This absence of cross-class interaction stems from sequence divergence at the inferred dimer interface residues responsible for coupling adjacent monomers within each Class I hexamer in structural models^15,45,46,65,66^. AlphaFold2 models of representative Class II and Class IV dimers indicated that the position and structure of this interface is preserved but composed of class-specific residues distinct from those found in Class I (Supplementary Fig.17).

We next compared conserved catalytic and regulatory motifs across eukaryotic PRPS classes (Supplementary Figs.18A-C). Catalytic residues required for the PRPP synthetase reaction were strictly conserved across all three classes, consistent with preservation of the core catalytic mechanism. Among the class-specific differences, changes within the ATP-recognition pocket were particularly notable because they involve the regulatory flexible (RF) loop, which contributes to both adenine recognition and allosteric regulation. In Class I, ATP is recognized within a dimeric pocket where the FLAG region from one subunit contributes E39 (*Homo sapiens* PRPS1 numbering), whose carboxylate side chain oxygen forms a hydrogen bond with the adenine N6 amino group, whereas the RF loop from the adjacent subunit contributes R96, whose guanidinium group forms a hydrogen bond with the adenine N1 atom (Supplementary Fig. 18D). Class IV largely retains the FLAG-region glutamate but harbors a conserved “STMER” motif within the RF loop that replaces the R96-equivalent residue with threonine. Because threonine lacks the long, positively charged guanidinium group of arginine, this substitution is expected to abolish the hydrogen bonding with adenine N1 while preserving the N6-contacting FLAG-region glutamate. Class II shows further divergence, containing a related “TTMER” RF-loop motif and lacking the conserved FLAG-region glutamate altogether, thus altering residues responsible for coordinating both the N1 and N6 positions of adenine. This loss of adenine-specific contacts is consistent with biochemical studies showing that plant Class II PRPS enzymes can use multiple diphosphoryl donors^17,18^. These changes suggest a stepwise relaxation of nucleotide-donor specificity, with Class IV representing an intermediate state between ATP-specific Class I and the broader nucleotide-donor utilization observed in Class II enzymes, providing a plausible evolutionary path toward biochemical innovation.

To experimentally test whether these motif-level differences correspond to measurable kinetic differences, we measured Class II and Class IV PRPS activity using a coupled assay. In this assay, PRPP generated from ATP and ribose-5-phosphate (R5P) by PRPS is sequentially converted by HPRT1 and IMPDH2 into IMP producing NADH, monitored by continuous fluorescence (Supplementary Figs.19A-D). Both recombinant Class II and Class IV PRPS catalyzed R5P, confirming that Class IV is an enzymatically active PRPS and demonstrating that metazoan Class II PRPS retains PRPP synthetase activity (Fig.3F). With ATP as the diphosphoryl donor, Class IV exhibited higher V_max_ and K_m_ values for R5P than Class II, indicating distinct turnover and substrate-affinity behavior between the two classes. While our analysis pinpointed and characterized the major evolutionary innovations, a complete structural and biochemical characterization of these enzymes – and a more robust comparison across all PRPS classes – will require future studies.

Collectively, the prokaryotic gene tree analysis (Fig. 2C), class-specific motif divergence (Supplementary Fig. 13), and class-level profile HMM scoring of ancestrally reconstructed bacterial PRPS sequences (Supplementary Figs. 20A and 20B) support the model summarized in Fig.3G: a stepwise bacterial diversification trajectory from a Class IA-like ancestor in which RF-loop remodeling and C-terminal changes (steps 1-2) yielded an early Class IV-like intermediate, bent dimer interface divergence (step 3) produced Class IV, and additional dimer interface and FLAG-region remodeling (step 4) generated Class II. The inclusion of LUCA in the model acknowledges that the precise ancestral state from which these classes derive cannot be unambiguously assigned from extant sequences alone. Class IV and Class II are thus not merely divergent sequence classes, but active, class-specific PRPS enzymes whose ancient evolutionary trajectory unfolded before or within bacteria prior to HGT to the stem eukaryote, preserving the core PRPS architecture while attaining distinct assembly and catalytic-regulatory states.

### Pervasive and recurrent pseudoenzymification across eukaryotic lineages and PRPS classes

Pseudoenzymes are catalytically inert homologs of active enzymes that have increasingly been recognized as prominent regulators of enzymes across diverse protein families^67–70^. Our previous work identified several pseudoenzymes^25^ that this study confirms were derived from Class I subclasses 1, 2, and 4. This prompted us to ask whether PRPS pseudoenzymification is a broader feature of eukaryotic PRPS evolution. By screening our expansive eukaryotic PRPS dataset for sequences possessing disruptive substitutions at residues required for catalysis we identified 37 independent pseudoenzymification events (Supplementary Table 4). The distribution of PRPS pseudoenzymes across eukaryotes revealed two broad trends (Fig.4A). First, pseudoenzymes were not restricted to a single PRPS class, with examples detected across multiple Class I and Class II subclasses, as well as in Class IV and cyano-PRPS. Second, pseudoenzymes occurred in representatives of all three major eukaryotic supergroups: Diaphoretickes, Amorphea, and Excavata. Thus, PRPS pseudoenzymification has occurred repeatedly across the eukaryotic PRPS repertoire. Compared with pseudoenzymification events reported across the 920 enzyme families catalogued in the Mechanism and Catalytic Site Atlas^71^, the PRPS enzyme family exceeds the highest reported counts, highlighting pseudoenzyme-based regulation of PRPS as a recurrent mechanism for tuning gene dosage, complex assembly, and flux control at a central metabolic chokepoint.

**Figure 4.**
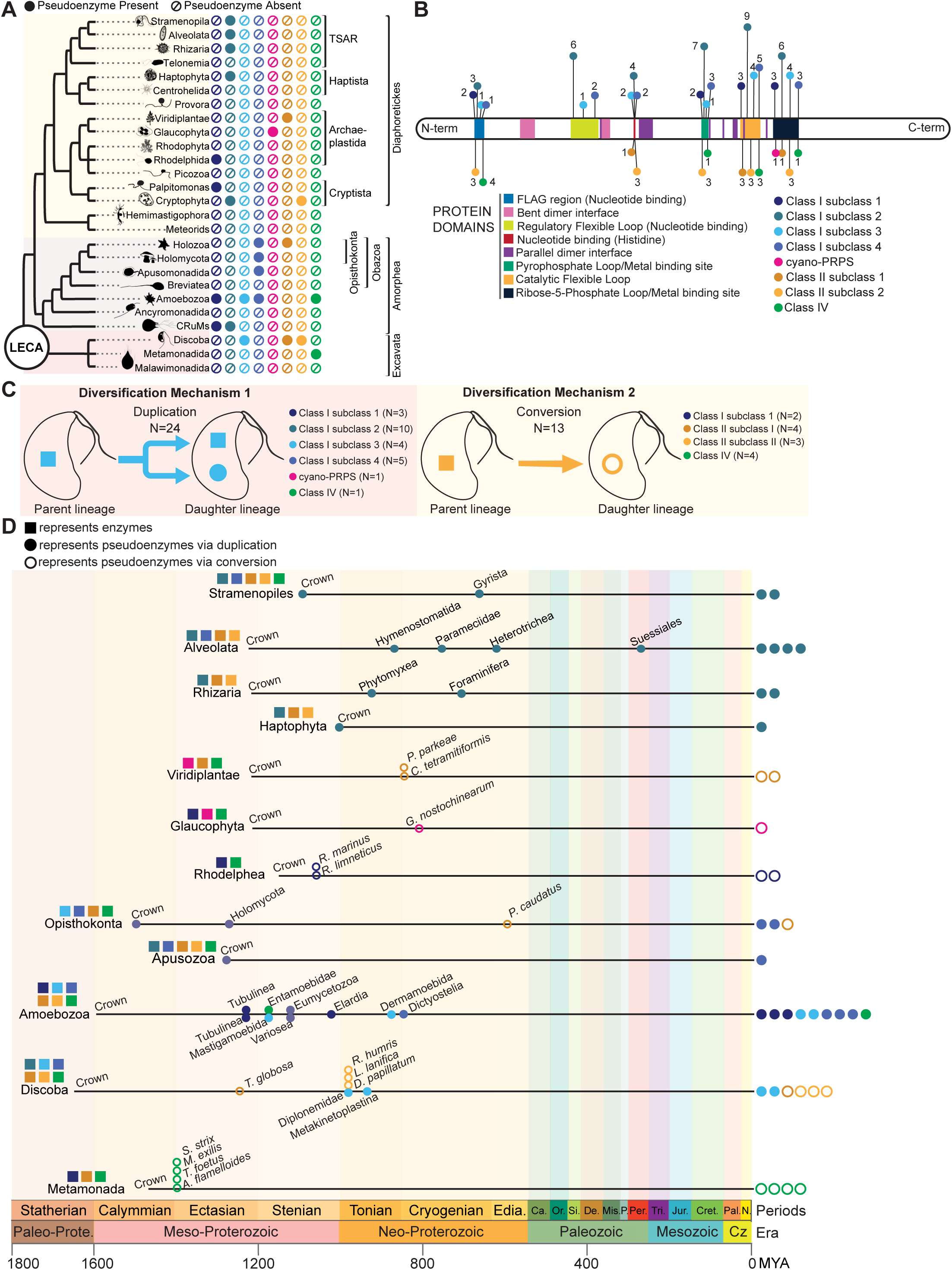
Pervasive and recurrent pseudoenzymification across eukaryotic lineages and PRPS classes. **(A)** Phylogenetic distribution of pseudoenzymes across PRPS classes and subclasses, showing their widespread presence across diverse eukaryotic lineages. **(B)** Structural and functional modifications in PRPS pseudoenzymes mapped along a generalized Class I-like PRPS polypeptide, with the number of altered features indicated. Modifications include changes in nucleotide-binding regions, catalytic and regulatory flexible loops, metal- and substrate-binding sites, and oligomerization interfaces. **(C)** Two primary mechanisms of pseudoenzymification are observed: gene duplication followed by divergence, and direct enzyme-to-pseudoenzyme conversion, with representative examples shown for each. To exclude species-specific duplications and lineage-restricted variants of unclear origin, only duplication events supported by at least two orthologs are recorded. **(D)** Timing of pseudoenzyme formation mapped onto a eukaryotic evolutionary timeline, indicating independent emergence across multiple lineages and geological intervals. Evolutionary timescale (millions of years ago, MYA) adapted from reference^27,29,72^. Symbols denote functional states as follows: squares, catalytically active enzymes; filled circles, pseudoenzymes arising from gene duplication; open circles, pseudoenzymes arising from direct conversion; and slashed circles, absence.

**Fig 5.**
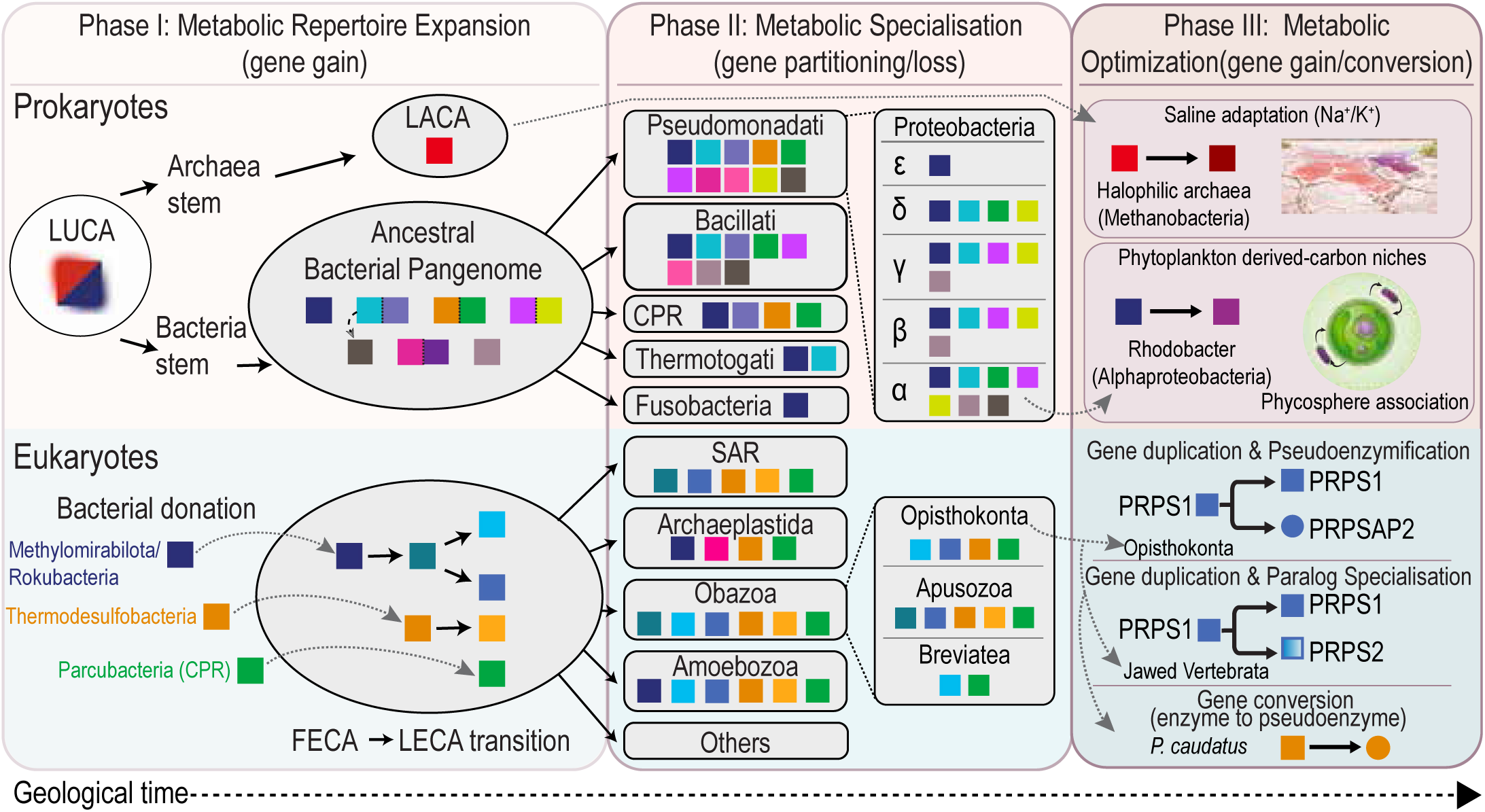
Macroevolutionary metabologenomic shift model of PRPS evolution. A three-phase macroevolutionary framework describing PRPS emergence and diversification across prokaryotic and eukaryotic lineages over geological time is shown. Colored squares denote distinct PRPS classes or homologs (see Figs.1 and 2 for annotation), and circles denote pseudoenzymes. In Phase I (metabolic repertoire expansion; gene gain), an expanded pool of PRPS homologs is generated. In Phase II (metabolic specialization; gene partitioning/loss), this expanded repertoire is pruned through lineage-specific retention and loss, reducing metabolic redundancy and contributing to genome streamlining. In Phase III (metabolic optimization; gene gain/conversion), the retained homologs are modified during ecological adaptation via duplication, specialization, and pseudoenzymification. In prokaryotes (top), early stem lineages (LUCA or the bacterial stem) establish metabolic autarky, involving generation of a diverse PRPS repertoire that typifies a metabolically expansive ancestral bacterial pangenome, whereas Archaea inherit a much more limited repertoire (Phase I). Partitioning of this repertoire across major and nested bacterial lineages accompanies metabolic specialization exemplified in the schema by allocation of PRPS among α, β, ψ, ο, and ε Proteobacteria from ancestral Pseudomonadati parent (Phase II). Further optimization produces lineage-specific innovations, including Class XII in halophilic archaea associated with saline adaptation and Class VI in Rhodobacter associated with phytoplankton-derived carbon niches (Phase III). In eukaryotes (bottom), PRPS expansion is initiated during the FECA-to-LECA transition via bacterial gene donation, involving contributions from multiple bacterial lineages. Candidate sources for major PRPS classes are supported by phylogenetic (gene tree), BLASTP, and HMM analyses of ancestral sequences. Establishment of a seven-gene PRPS repertoire in LECA occurs through several gene duplication events (Phase I). This repertoire is subsequently partitioned by hierarchical retention and loss across supergroups and daughter lineages, exemplified in the schema by PRPS gene losses from the present Obazoan ancestor to the Opisthokonta ancestor (Phase II). Subsequent innovation arises through gene duplication and pseudoenzymification (illustrated showing PRPSAP2 from PRPS1 in Opisthokonta), gene duplication and paralog specialization (illustrated showing PRPS2 from PRPS1 in Jawed Vertebrata), and direct enzyme-to-pseudoenzyme conversion (illustrated showing Class II subclass 1 in *P. caudatus*) (Phase III). Together, these patterns support a unified macroevolutionary model in which metabolic innovation proceeds through sequential phases of expansion, lineage-specific pruning, and functional optimization over deep evolutionary time.

To examine how PRPS pseudoenzymification is coupled to neofunctionalization, we mapped disruptive modifications across PRPS pseudoenzymes (Fig.4B and Supplementary Table 4). As expected, pseudoenzymes carried substitutions at residues required for catalytic activity, but these changes were typically accompanied by additional modifications across nucleotide-binding regions, regulatory and catalytic flexible loops, metal- and substrate-coordinating residues, and oligomerization interfaces. Individual pseudoenzymes carried different combinations of these alterations: although certain critical residues were recurrently disrupted, no single uniform pattern was shared across all pseudoenzymes. This heterogeneity suggests that PRPS pseudoenzymification does not follow one fixed molecular route, but instead generates variants predicted to affect distinct biochemical or assembly-related properties. Because core intersubunit interfaces were largely preserved across PRPS pseudoenzymes, they are likely to associate with the retained active PRPS class from which they emerged. Therefore, rather than acquiring moonlighting functions, pseudoenzymes primarily optimize PRPS complex assembly and activity as shown previously^25^. We propose that the regulatory outcome of pseudoenzymification is shaped by the specific sites that are modified, determining whether a pseudoenzyme influences assembly/dynamics, substrate recognition, catalytic coordination, and/or allostery of the active PRPS enzyme it engages.

We next explored how PRPS pseudoenzymes arose. Two major routes were evident (Fig.4C). The more frequent route involved gene duplication followed by divergence, producing a catalytically modified paralog alongside a retained active copy in the same lineage. The other route involved direct conversion of an existing PRPS gene into a pseudoenzyme-like form, with no retained duplicate. Critically, direct conversion was largely restricted to species already encoding another active PRPS from the same class indicating that the two routes operate under distinct genomic prerequisites: duplication frees a daughter to diverge while the parent maintains catalysis, whereas direct conversion can only proceed where a pre-existing backup preserves catalytic flux, permitting the converted gene to acquire a non-catalytic regulatory role whether for recalibrating gene-dosage or fine-tuning PRPS complex activity.

Mapped onto a eukaryotic evolutionary timeline^29,72^, these 37 independent pseudoenzymification events span deep crown-group nodes including the ancestors of Opisthokonta and Stramenopila to more recent species-restricted innovations such as the Class II subclass 1 conversion in *Priapulus caudatus* (Fig.4D). Their recurrence across distant taxa and disparate timescales argues against a single ancient burst and indicates that no PRPS class was uniquely predisposed to catalytic inactivation. Together with the preservation of assembly-associated interfaces and the two routes of pseudoenzyme emergence, these patterns establish pseudoenzymification alongside gene duplication, partitioning, and specialization as an additional evolutionary mechanism that has optimized the eukaryotic PRPS repertoire for metabolic flexibility.

## DISCUSSION

Here, we have elucidated the evolutionary arc of a universal enzyme that predates the LUCA – the phosphoribosyl pyrophosphate synthetase – to reveal how a comprehensive array of protein innovation and diversification mechanisms have collectively operated across deep time to shape the panoply of PRPS enzyme forms that exist today. A key finding is that multiple PRPS gene duplications occurred independently in the stem lineages of both bacteria and eukaryotes. Indeed, the last bacterial common ancestor (LBCA) and LECA each harbored more PRPS paralogs than any of their progeny, which likely reflects the nature of the selective pressures that existed before and after their formation. Mechanistically, we propose a macroevolutionary shift model (Fig.5) whereby a biomolecule-scarce early Earth habitat necessitated expansive PRPS repertoires for metabolic self-sufficiency and supreme adaptability. As symbiont-rich ecological niches replete with biomolecules developed, a transition occurred from the ancestral slowly proliferating organisms with bulky genomes to more metabolically specialized and genomically streamlined taxa encoding fewer PRPS genes. Later, as competition for ecological niches and biomolecular resources between organisms intensified, bacteria (and in one instance, Archaea) adapted by creating new PRPS enzyme classes, whereas eukaryotic organisms created PRPS-associated pseudoenzymes to fine-tune isozyme regulation.

Beyond using PRPS evolution to reveal macroevolutionary phenomena, we introduce a robust approach for chronological ordering of gene duplications within the first eukaryotic common ancestor (FECA)-to-LECA stem lineage, and we demonstrate its utility with two high-confidence proof-of-concept examples (Supplementary Figs.6-8). In addition, although the precise horizontally transferred genes from bacteria to the FECA around 3.5 billion years ago remain unspecified, our results provide compelling evidence that acquisition of the three PRPS genes ranks among the earliest horizontal transfer events in eukaryogenesis. First, the total absence of Archaea-derived PRPS in eukaryotes suggests a fundamental reliance on bacterially donated PRPS prior to LECA’s radiation. Second, the extensive paralog expansion (three class I and one class II duplication) occurred entirely within the FECA-to-LECA stem. Most importantly, the unique distribution of splice sites – consistent within classes I, II, and IV but distinct between them – indicates that these genes were acquired prior to the spread of group II introns and/or functional spliceosomal machinery. The retention of class-specific splice-site positions in PRPS paralogs indicates these gene duplications occurred after the origin of split-genes in early eukaryotes – an event commonly attributed to endosymbiosis of the protomitochondrion^73^, although recent studies also support possible archaeal-host^74^ or chimeric contributions^75^. Though the bacterial donor lineages of the three ancestral eukaryotic PRPS genes remain unclear, our probabilistic approach disfavors an alphaproteobacterial source (and other less likely lineages), narrowing the search space for future phylogenomic studies. Extending our methodology to additional phosphoribosyltransferase family members found in eukaryotes could reveal co-evolutionary patterns and ancient operons, thereby providing a direct path to identifying and reconstructing the specific bacterial lineages that were the first participants in early eukaryogenesis. Our study also defines seven well-characterized genes that do not co-segregate in eukaryotes, which can serve as powerful genomic landmarks for future synteny analysis in the reconstruction of LECA’s genome or its partitioning during eukaryotic diversification.

Finally, by mapping and re-tracing the PRPS protein sequence space across the breadth of biological history, we establish a roadmap for downstream structure-function investigations geared toward the rational design of metabolic or therapeutic innovations. More consequentially, this evolution-grounded framework serves as a blueprint for deciphering how evolutionary ‘tinkering’ drives the emergence of functional complexity – a methodology that can be applied to any individual protein and eventually scaled across protein families to reveal broader principles governing molecular innovation.

## METHODS

### Dataset construction

Amino acid sequences from well-annotated PRPS enzymes in representative model organisms were used as queries to identify homologous sequences in the NCBI non-redundant database^76^ using BLASTP^77^. Candidate sequences were curated based on sequence similarity, domain architecture, and the presence of universally conserved PRPS residues. To expand taxonomic coverage, additional sequences were derived from publicly available transcriptomic datasets obtained from the Sequence Read Archive (SRA)^78^. Datasets were selected based on sequencing quality and availability of associated metadata. Transcript assemblies were mined using tBLASTn against curated PRPS reference sequences to identify candidate homologs. Putative open reading frames (ORFs) were identified from transcript sequences, and corresponding protein sequences were inferred based on canonical start and stop codons. Predicted proteins were retained if they exhibited significant similarity to known PRPS sequences and contained characteristic conserved residues. Taxonomic annotations for most sequences were obtained from the NCBI Taxonomy database.

### Splice site analysis

For splice site analysis, genomic DNA or mRNA sequences corresponding to each PRPS homolog were retrieved from NCBI^76^ and Ensembl^79^ databases. In cases where exon-intron annotations were not explicitly available, homologous sequences were used to infer coding regions, and exon-intron boundaries were assigned based on canonical (GT/AG) and non-canonical (GC/AG) splice junction rules. Predicted splice junctions were further evaluated against available transcriptomic data from the same species or closely related organisms when possible. All exonic and intronic sequences used in this analysis are available via Figshare.

### Phylogenetic tree construction

Representative PRPS protein sequences were aligned using MAFFT (L-INS-i). Partial sequences and those exhibiting long-branch artifacts were removed prior to downstream analysis. Poorly aligned regions were trimmed using ClipKit in smart-gap mode. Maximum likelihood phylogenetic analysis was performed using IQ-TREE v3.0.1^80^. The optimal substitution model was selected using ModelFinder from a predefined set of substitution matrices (LG, WAG, VT, JTT, Blosum62, DCMut, Dayhoff, and Poisson). Model selection was based on the best Bayesian Information Criterion (BIC) score. Tree inference was conducted under the selected model with automatic thread detection and a fixed random seed. Branch support was evaluated using ultrafast bootstrap approximation (1,000 replicates) with nearest-neighbor interchange (NNI) optimization applied during bootstrap analysis. Final trees were visualized and annotated using iTOL (version 7.5.1)^81^.

### Ancestral sequence reconstruction

Ancestral sequence reconstruction (ASR) of PRPS enzymes was performed using a two-phase maximum-likelihood framework to establish rooted topologies while minimizing outgroup-driven bias. For eukaryotic ASR, orthologous PRPS sequences were curated across major lineages for the specified class, with pseudoenzymes and duplicated paralogs removed to retain the least diverged ortholog per species. For bacterial Class II and Class IV ASR, most bacterial PRPS sequences were used except for highly divergent sequences. Sequences were aligned using MAFFT (L-INS-i) and trimmed to remove poorly aligned regions based on gap-fraction thresholds (terminal 0.10; internal 0.50). Phylogenetic inference for eukaryotic ASR was performed using IQ-TREE v3.0.1 under hierarchical topological constraints reflecting established deep eukaryotic relationships, including major supergroups (Diaphoretickes, Amorphea, and Excavata) and nested lineages such as SAR, Archaeplastida, and Discoba. Bacterial ASR was performed without hierarchical constraints. In Phase A, the best-fitting substitution model was selected using ModelFinder, followed by tree inference and ASR in the presence of outgroups. Branch support for bacterial ASR trees was assessed using ultrafast bootstrap approximation (1,000 replicates) and SH-like approximate likelihood ratio tests (1,000 replicates), with nearest-neighbor interchange (NNI) optimization applied during bootstrapping. In Phase B, outgroups were removed and replaced with a minimally informative “ghost” sequence grafted at the ingroup MRCA. The ghost sequence encoded residues at a small number of strictly invariant ingroup positions (typically five) and was otherwise ambiguous. ASR was then repeated on the fixed Phase A topology to obtain final ancestral sequences.

### Computing protein embeddings from ESM-2

PRPS protein sequences from eukaryotic and prokaryotic organisms were embedded using the pretrained transformer-based protein language model ESM-2 (esm2_t33_650M_UR50D)^8^. Sequence sets included representative homologs from bacteria, archaea, and eukaryotes, as well as ASR of eukaryotic PRPS classes. For each sequence, a fixed-length representation was obtained by mean pooling residue embeddings from the final transformer layer, with model weights held fixed and gradient computation disabled, yielding a 1280-dimensional embedding vector per protein. Low-dimensional projections of embedding space were generated using Uniform Manifold Approximation and Projection (UMAP)^82^. Dimensionality reduction was performed using cosine distance, with parameters n_neighbors = 15 and min_dist = 0.1 for eukaryotic datasets, and n_neighbors = 30 and min_dist = 0.05 for prokaryotic datasets, using a fixed random seed.

### Distance-based protein clustering

Eukaryotic PRPS protein sequences were aligned using MAFFT^83^ (L-INS-i) and subsequently trimmed to remove columns with a gap-fraction threshold of 0.5. Partial sequences and sequences exhibiting long-branch artifacts were removed prior to analysis. A gap-aware pairwise p-distance matrix was computed from the trimmed alignment using only retained columns and normalized by the number of comparable (non-gap) sites for each sequence pair. Unsupervised clustering was performed using spectral clustering across a range of cluster numbers (k = 2-9), and clustering quality was evaluated using silhouette scores. Nonlinear dimensionality reduction was performed using UMAP with the pairwise distance matrix supplied as a precomputed metric. Three-dimensional embeddings were generated using n_neighbors = 15, min_dist = 0.2, and a fixed random seed.

### BLASTP analysis of ancestral PRPS sequences

Pairwise sequence similarity between ASR of eukaryotic PRPS sequences and prokaryotic homologs was assessed using BLASTP^77^. Each query was searched against a curated database of 33,248 PRPS sequences (28,560 bacterial and 4,688 archaeal). A local BLAST database was constructed using the BLAST+ suite (makeblastdb), and searches were performed using blastp with parameters corresponding to NCBI BLAST defaults (BLOSUM62 substitution matrix, gap opening penalty 11, gap extension penalty 1, word size 3, and composition-based statistics enabled). Low-complexity filtering was disabled (-seg no).

### HMM profiling of ancestral PRPS sequences

Profile hidden Markov models (HMMs)^61^ were constructed for representative bacterial and archaeal PRPS classes using hmmbuild (HMMER v3)^84^. Multiple sequence alignments for each class were generated using MAFFT (L-INS-i), and resulting profiles were combined into a searchable database using hmmpress. Query sequences included ASR of PRPS (LECA Class I, Class II, Class IV, and ACD-PRPS). Each query was searched against the HMM panel using hmmsearch, and bit scores were extracted for all query-model comparisons. For each query, the top-scoring HMM and the difference in bit score relative to the next-best model (Δ score) were calculated. Analyses were repeated across multiple alignment strategies (MAFFT, Clustal Omega, MUSCLE) to assess robustness of score distributions.

### Diagnostic residue analysis

PRPS protein sequences were grouped into predefined classes or subclasses based on protein embeddings, distance-clustering and phylogenetic analysis. Multiple sequence alignments were generated using MAFFT (L-INS-i). For large datasets, a class-balanced subset was first aligned to generate a backbone alignment, and remaining sequences were incorporated using the --addfull option. Diagnostic residues were identified by calculating amino acid frequencies at each alignment position within and across groups. For each residue, within-group frequency was computed over non-gap positions, and coverage was defined as the fraction of non-gap residues within (coverage-in) and outside (coverage-out) the focal group. Residues were classified as diagnostic if they satisfied the following criteria: frequency-in ≥ 0.65, Δmax ≥ 0.50, coverage-in ≥ 0.75, coverage-out ≥ 0.60, and statistical significance by one-sided Fisher’s exact test with Benjamini-Hochberg correction (q ≤ 0.01), where Δmax is the difference between the in-group residue frequency and the maximum frequency observed in any other group. Where multiple residues satisfied these criteria at a given position, the highest-ranking residue was retained based on Δmax, followed by within-group frequency and log₂ enrichment. Diagnostic residues were summarized as group-by-position matrices and visualized as heatmaps using normalized residue frequencies.

### Sequence logo generation

To visualize residue conservation across the PRPS family, sequence logos were generated from trimmed multiple sequence alignments using WebLogo^85^. Eukaryotic alignments were trimmed using a gap-fraction threshold of 0.3, whereas prokaryotic full-length alignments were filtered using a gap-fraction threshold of 0.1. Sequence logos were generated independently for each class from the resulting filtered alignments.

### Cell culture

HEK293T cells were purchased from American Type Culture Collection (ATCC). Cells were maintained in Dulbecco’s modified Eagle medium (DMEM) containing 10% (v/v) fetal bovine serum (FBS) and 1x penicillin/streptomycin.

### Plasmids and transfection

cDNA sequences encoding *Corallochytrium limacisporum* Class IV PRPS (annotated from SRX738098 and SRX732498) and *Branchiostoma lanceolatum* Class II PRPS (GenBank ID: CAH1247366.1) were designed and synthesized as gene fragments (Twist Bioscience). These fragments were cloned into FUGW-blasticidin (as described in reference^25^) and pET21a-6×His vector backbones using HiFi DNA Assembly (New England Biolabs #E2621L) to generate mammalian and bacterial expression constructs, respectively. For human HPRT1, the coding sequence was amplified from cDNA derived from 786-O cells (as described in Bischoff et al.) and cloned into the pET21a-6×His vector using HiFi DNA Assembly. HEK293T cells were transfected using Lipofectamine 3000 (Invitrogen #L3000001) according to the manufacturer’s instructions.

### Recombinant protein purification

*Branchiostoma lanceolatum* Class II PRPS, *Corallochytrium limacisporum* Class IV PRPS, and human hypoxanthine phosphoribosyltransferase (HPRT1) were expressed in *E. coli* BL21-DE3 cells (New England Biolabs #C2527I) transformed with pET21a vectors encoding N-terminal 6×His-tagged proteins. For each construct, a 2 mL overnight LB-ampicillin culture was used to inoculate 1 L LB-ampicillin medium (1:500, v/v) and grown at 37°C with shaking to an OD₆₀₀ of ∼0.8. Protein expression was induced with 1 mM IPTG (Thermo #R0392), followed by incubation at 30°C overnight. Cells were harvested by centrifugation (3,500 × g, 20 min, 4 °C) and stored at -80°C prior to purification. Cell pellets were resuspended in 50 mL lysis buffer and lysed by sonication on ice. For Class II and Class IV PRPS, the lysis buffer contained 25 mM Tris-Cl pH 8.0, 100 mM NaCl, 0.1 mM EDTA, 0.1% Triton X-100, 5% glycerol, 5 mM MgCl₂, 2.5 mM DTT, 0.5 mg/mL lysozyme (Thermo #89833), protease inhibitor cocktail (Pierce #A32963), and benzonase (100 U/mL; Millipore #E1014). For HPRT1, the lysis buffer consisted of 10 mM Tris-Cl pH 8.0, 100 mM NaCl, 0.1 mM EDTA, 0.1% Triton X-100, 5 mM MgCl₂, 0.5 mg/mL lysozyme, protease inhibitors, and benzonase. Lysates were clarified by centrifugation (20,000 × g, 30 min, 4 °C), supplemented with 30 mM imidazole, and filtered through 0.22 μm membranes. Clarified lysates were loaded onto a pre-equilibrated 5 mL HisTrap HP Ni²⁺ affinity column (Cytiva #17524802). Columns were washed with binding buffer (20 mM sodium phosphate, 500 mM NaCl, 30 mM imidazole, pH 8.0), and proteins were eluted using the same buffer containing 500 mM imidazole. Eluted proteins were concentrated using 10 kDa MWCO centrifugal concentrators (Thermo #88528) and further purified by size-exclusion chromatography on a Superose 6 Increase 10/300 column (ÄKTA Pure system). For Class II and Class IV PRPS, the gel filtration buffer consisted of 50 mM HEPES pH 8.0, 400 mM NaCl, 1 mM DTT, and 5% glycerol. For HPRT1, the buffer contained 50 mM Tris-Cl pH 8.4, 300 mM NaCl, 1 mM TCEP, and 5% glycerol. Peak fractions were pooled, analyzed by SDS-PAGE and Coomassie staining, snap-frozen in liquid nitrogen, and stored at -80 °C. For analytical size-exclusion chromatography, aliquots corresponding to the major peak fractions from Superose 6 runs were run on a pre-equilibrated Yarra 3 μm SEC-2000 column (100 mM NaH₂PO₄, pH 7.0, 100 mM NaCl) using a Vanquish UHPLC system (Thermo Scientific). The column has an optimal fractionation range of ∼1-300 kDa. Gel filtration calibration kits (GE #28-4038-41, GE #28-4038-42, Sigma #MWGF200-1KT) were used as molecular weight standards. Chromatograms were processed using Unicorn (Cytiva Life Sciences) and Chromeleon (Thermo Scientific) software and plotted in GraphPad Prism 8.

### AlphaFold2 structure prediction

AlphaFold2^86^ modellings were generated using the Alphafold2_mmseq2^87^ notebook executed on Google Collaboratory cloud^88^. Predicted models were ranked (1-5) based on per-residue Local Distance Difference Test (pLDDT) scores, which provide a confidence estimate for each residue on a scale of 0-100. The model with the highest average pLDDT score was selected for downstream structural analysis. Visualization and inspection of predicted structures were performed using ChimeraX^89^.

### SDS-PAGE and Western blotting

Cells were rinsed once with ice-cold PBS and lysed in RIPA buffer (Thermo Scientific #89901) supplemented with 1× protease and phosphatase inhibitor cocktail (Thermo #78446). Lysates were clarified by centrifugation and mixed with 1× Laemmli sample buffer prior to electrophoresis on 10% TGX FastCast gels (Bio-Rad #1610173). Proteins were then blotted onto 0.2 μm PVDF membranes using a Trans-Blot Turbo system (Bio-Rad). Membranes were blocked for 40 min at room temperature in 5% (w/v) nonfat milk prepared in Tris-buffered saline containing 0.1% Tween-20 (TBS-T). Following blocking, membranes were incubated overnight at 4°C with primary antibodies diluted 1:1000 in 3% BSA in TBS-T. Primary antibodies included PRPS1/2 (Santa Cruz #sc-100822), PRPS1/2/3 (Santa Cruz #sc-376440), PRPSAP1 (Santa Cruz #sc-398422), PRPSAP2 (Proteintech #17814-1-AP), and ALFA-HRP (Synaptic Systems #N1505-HRP). After washing, membranes were incubated with HRP-conjugated secondary antibodies (Jackson ImmunoResearch; 1:25,000 dilution in 5% milk in TBS-T), including anti-mouse (#115-035-003) and anti-rabbit (#111-035-003). Signal detection was performed using chemiluminescent substrates (Thermo Scientific) and imaged on a ChemiDoc system (Bio-Rad #12003153). Images were processed using Image Lab software (v5.2.1, Bio-Rad). For re-probing, membranes were stripped using Restore PLUS Western blot stripping buffer (Thermo #46430). For Coomassie staining, QC Colloidal Coomassie stain (Bio-Rad #161-0803) was used according to the manufacturer’s instructions.

### Analytical SEC from mammalian cells

HEK293T cells were lysed in non-denaturing lysis buffer (50 mM Tris-Cl, pH 7.5, 200 mM NaCl, 1% digitonin, 1 mM TCEP, 1 mM MgCl₂, benzonase, and 1× protease and phosphatase inhibitor cocktail) for 20 min on ice. Lysates were clarified by centrifugation (15,000 × g, 15 min, 4 °C) and filtered through a 0.22 μm membrane. 200 μg of total protein was loaded onto a pre-equilibrated Superose 6 Increase 3.2/300 column (100 mM NaH₂PO₄ pH 7.0, 100 mM NaCl) and separated using a Vanquish UHPLC system (Thermo Scientific) according to the manufacturer’s instructions. Following elution past the void volume, fractions were collected, concentrated using a 3 kDa MWCO centrifugal filter (Thermo #88512), and analyzed by Western blotting. Internal molecular weight standards used for calibration are described in reference^25^.

### Co-immunoprecipitation analysis

HEK293T cells transiently overexpressing C-terminally ALFA tagged^90^ Class II PRPS or Class IV PRPS were lysed in non-denaturing lysis buffer (50 mM Tris-Cl pH 7.5, 200 mM NaCl, 1% digitonin, 1 mM TCEP, 1 mM MgCl₂, benzonase, and 1× protease and phosphatase inhibitor cocktail) for 20 min on ice. Lysates were clarified by centrifugation (15,000 × g, 15 min, 4°C) and incubated with equilibrated anti-ALFA nanobody-conjugated agarose beads (ALFA Selector ST, NanoTag #N1511) for 1 h at 4°C with gentle rotation. Following incubation, the supernatant was removed, and the beads were washed four times with wash buffer (25 mM Tris-Cl pH 7.5, 0.5 M NaCl, 0.5% Triton X-100). Bound proteins were eluted by boiling the beads in 2× Laemmli sample buffer at 95°C for 10 min prior to SDS-PAGE analysis.

### PRPS activity assay

Reactions were performed in black 96-well plates in a final volume of 100 μL containing 1× activity buffer (50 mM Tris-Cl, 50 mM sodium phosphate, 10 mM MgCl₂, 1 mM DTT, 0.01% Triton X-100, pH 7.5), 1 mM ATP, 1 mM NAD, and 1 mM hypoxanthine, supplemented with HPRT1 (0.2 μM) and IMPDH2-Y12A (0.2 μM), and PRPS at the indicated concentrations. Ribose-5-phosphate (R5P) was added at varying concentrations from concentrated stock solutions. Following brief mixing and incubation at 37°C, NADH production was monitored by continuous fluorescence (excitation 335 nm, emission 440 nm) at regular intervals for up to 2 h using a microplate reader. An NADH standard curve prepared in parallel was used to convert fluorescence units to NADH concentrations. Initial rates were determined from the linear portion of the progress curves and normalized to the amount of PRPS enzyme. Kinetic parameters were obtained by fitting the data to the Michaelis-Menten equation. IMPDH2 constructs and protein were a gift from the Kollman lab and the protein purification is described in reference^91^.

### Statistics and reproducibility

Diagnostic residue enrichment was assessed using custom Python scripts, with one-sided Fisher’s exact tests performed using SciPy followed by Benjamini–Hochberg correction for multiple testing. GraphPad Prism was used generating kinetic parameters shown in Figure 4. Sample sizes, replicates, and statistical tests used are indicated in each figure legend.

## DATA AVAILABILITY

All data supporting the findings of this study are available within the Article and its Supplementary Information (Note: Additional data deposited in a public repository (Figshare) will be available upon request and fully released during publication). All sequence datasets used in this study, including curated PRPS homologs from bacteria, archaea, and eukaryotes, are provided in Supplementary Tables 1-4 and will be available in FASTA format via Figshare. Publicly available datasets include sequence data from NCBI and Ensembl genome resources, as well as transcriptomic datasets from the Sequence Read Archive (SRA). Accession numbers for representative sequences are provided in the Supplementary Tables. The Figshare repository will contain all processed datasets generated in this study, including sequence alignments, phylogenetic trees, ancestral sequence reconstructions, protein embedding data, diagnostic residue matrices, and BLASTP and profile HMM analysis results. All unique reagents generated in this study will be made available by the corresponding author upon reasonable request and completion of a Materials Transfer Agreement.

## CODE AVAILABILITY

All scripts used for sequence processing, protein embedding, phylogenetic analysis, ASR, clustering, and visualization will be available upon request and released during publication.

## ACKNOWLEDGEMENTS

This work received support by NIH grant (R35GM133561) to J.T.C. and NIH UL1TR001425, 1R01AG083628 and R01CA287260 to J.M. J.T.C. is recipient of LCRF Leading Edge Award. We thank the Kollman group at the University of Washington (J. Kollman, A. O’Neill, and K. Hvorecny) for providing IMPDH2 constructs and protein, as well as thank K. Hvorecny for discussions. The silhouette images in phylogenetic trees were downloaded from Phylopic (http://phylopic.org/) or designed using Adobe Illustrator (version 30.4). All images downloaded were freely available for reuse under a Public Domain license. Schema generation and figure formatting was done using Adobe Illustrator (version 30.4). Phylogenetic trees were visualized and annotated using iTOL (version 7.5.1).

## AUTHOR CONTRIBUTIONS

Conceptualization, B.R.K. and J.T.C; Methodology, B.R.K., J.M., J.T.C.; Investigation, B.R.K. and J.T.C; Formal analysis, B.R.K., and J.T.C; Writing – Original Draft, B.R.K. and J.T.C.; Writing – Review & Editing, B.R.K., J.M., J.T.C; Visualization, B.R.K. and J.T.C.; Project administration, J.T.C.; Supervision, J.M. and J.T.C.; Funding Acquisition, J.T.C.

## ETHICS DECLARATION

### Competing Interests

B.R.K. and J.T.C. have filed patent applications on this work. The remaining authors declare no competing interests.

### Declaration of generative AI and AI-assisted technologies

During the preparation of this work, the authors used ChatGPT to assist with aspects of computational workflow development (including phylogenetic and ancestral sequence reconstruction analyses) and with minor language editing (grammar and syntax). All outputs generated by this tool, including code, commands, and text, were reviewed, validated, and revised by the authors as necessary. The authors take full responsibility for the content, accuracy, and integrity of this publication.

---

**Supplementary Figure 1.**
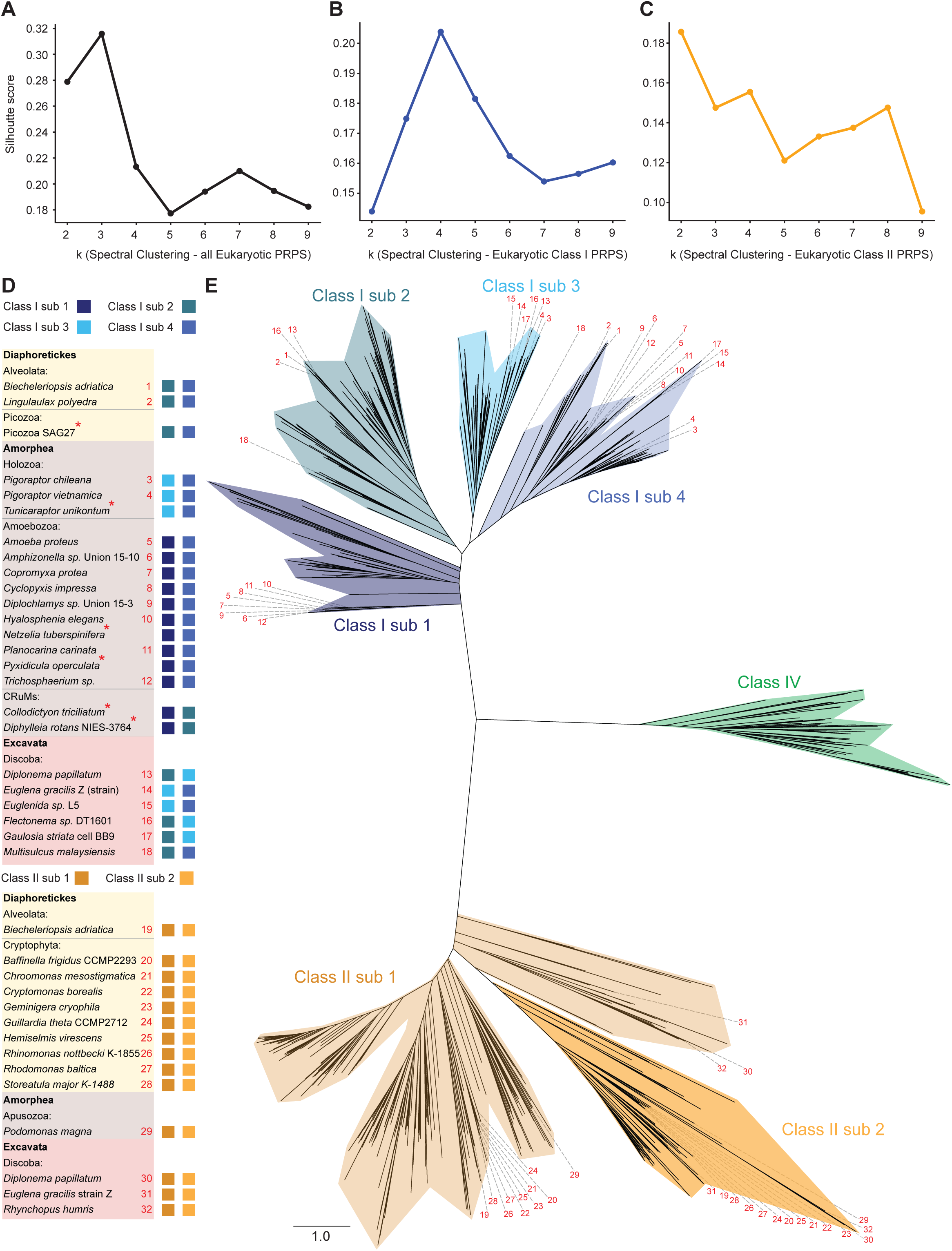
Robust clustering and subclass co-existence validate duplications of ancestral eukaryotic PRPS genes. **(A-C)** Silhouette score analysis of spectral clustering on a pairwise distance matrix of all eukaryotic PRPS sequences (A), eukaryotic Class I PRPS sequences (B), and eukaryotic Class II PRPS sequences (C) across k = 2-9. **(D)** Distribution of PRPS subclasses across eukaryotic lineages. Presence of subclasses is shown across major eukaryotic supergroups, highlighting species encoding multiple PRPS subclasses. Asterisks denote species for which one or more PRPS sequences were partial and excluded from the gene tree analysis in Figure 1C. **(E)** Positions of PRPS homologs from multi-subclass species across the inferred eukaryotic PRPS gene tree, with numbered labels corresponding to species shown in (D).

**Supplementary Figure 2.**
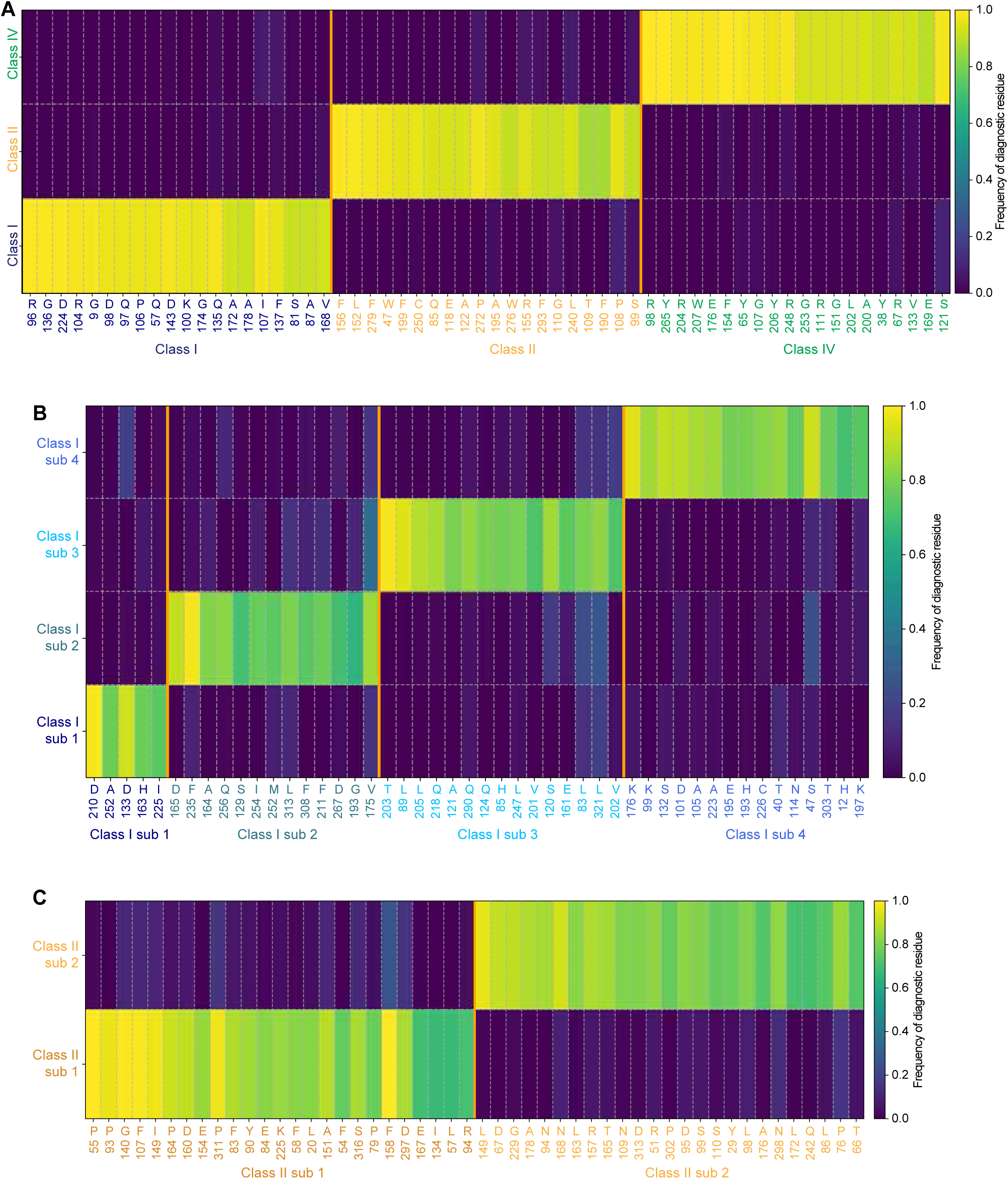
Diagnostic residue signatures define distinct PRPS classes and subclasses in eukaryotes. **(A-C)** Heatmaps showing diagnostic residues distinguishing eukaryotic PRPS classes (A), eukaryotic Class I subclasses (B), and eukaryotic Class II subclasses (C). Rows correspond to classes or subclasses, and columns represent alignment positions meeting stringent diagnostic criteria (frequency, coverage, and statistical criteria – see Methods). Cells indicate residue frequency (0-1) within each group, calculated from non-gap residues at each position. Only the top diagnostic sites per class are shown. Diagnostic site positions for Class I, Class II, and Class IV PRPS shown in (A), for Class I subclasses shown in (B), and for Class II subclasses shown in (C) correspond to representative organism sequences shown in Supplementary Figure 3. Group sizes for (A): Class I (n = 417), Class II (n = 422), Class IV (n = 68). Group sizes for (B): Class I sub 1 (n = 71), sub 2 (n = 155), sub 3 (n = 84), sub 4 (n = 107). Group sizes for (C): Class II sub 1 (n = 341), sub 2 (n = 81).

**Supplementary Figure 3.**
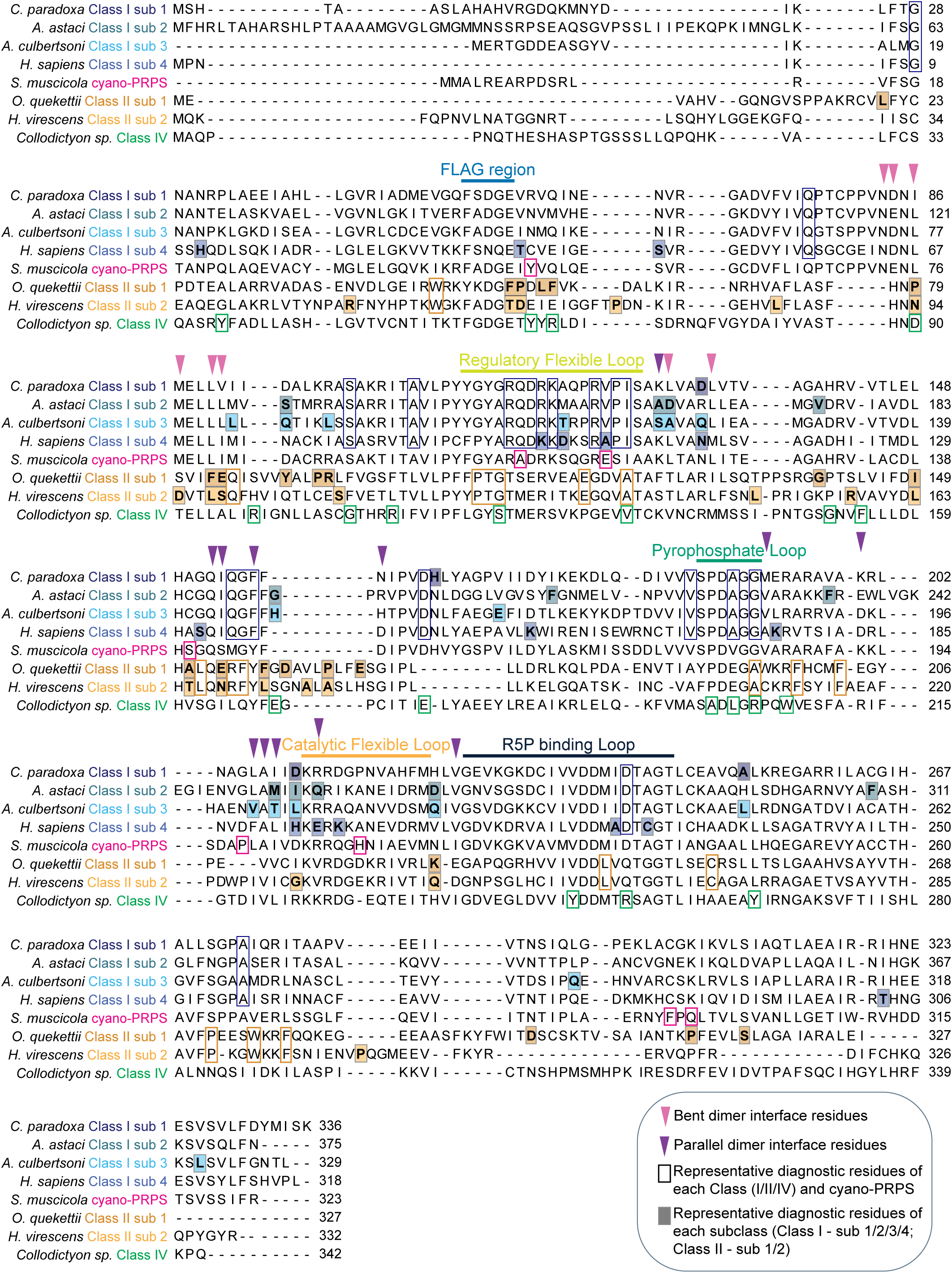
Diagnostic residues map to both established functional regions and residues of undetermined significance across PRPS classes/subclasses. Aligned PRPS sequences from representative eukaryotic species across classes and subclasses, with diagnostic residues highlighted. Functional regions, including the FLAG region, regulatory flexible loop, pyrophosphate loop, catalytic flexible loop, and R5P-binding loop, are shown. Residues contributing to parallel and bent dimer interfaces are indicated. Diagnostic residues correspond to class-level (Class I, II, IV, and cyano-PRPS) and subclass-level signatures shown in Supplementary Figure 2.

**Supplementary Figure 4.**
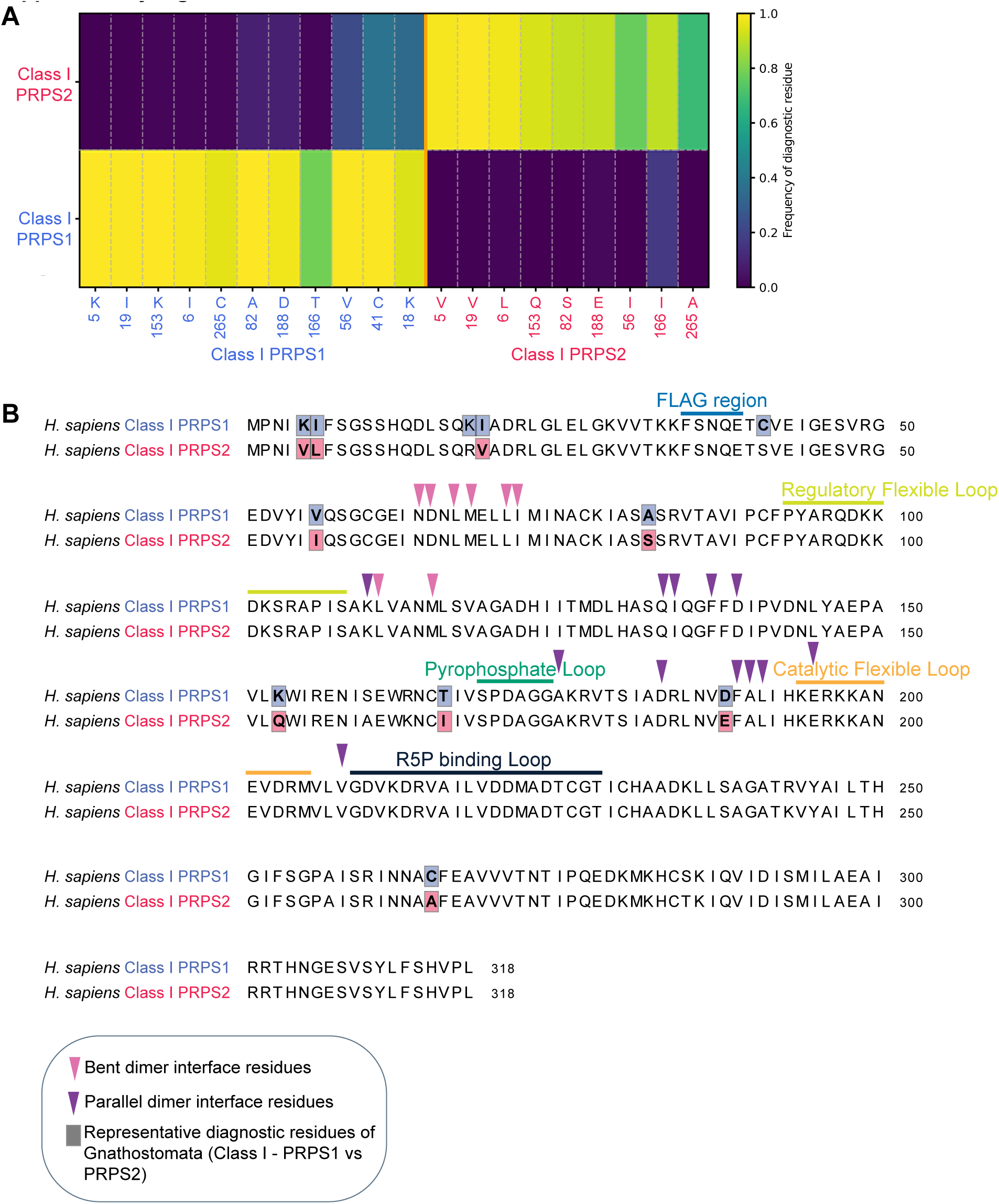
Diagnostic residues distinguish Gnathostomata PRPS1 and PRPS2. **(A)** Heatmap of diagnostic residues distinguishing PRPS1 and PRPS2. Rows correspond to PRPS1 and PRPS2, and columns represent alignment positions meeting stringent diagnostic criteria (frequency, coverage, and statistical criteria – see Methods). Cells indicate residue frequency (0-1) within each group, calculated from non-gap residues at each position. Diagnostic site positions for PRPS1 and PRPS2 shown correspond to human PRPS1 and PRPS2, respectively shown in (B). Group sizes: PRPS1 = 150, PRPS2 = 150. **(B)** Representative sequence alignment of human PRPS1 and PRPS2 with highlighted diagnostic residues identified in (A). Functional regions, including the FLAG region, regulatory flexible loop, pyrophosphate loop, catalytic flexible loop, and R5P-binding loop, are shown. Residues contributing to parallel and bent dimer interfaces are indicated.

**Supplementary Figure 5.**
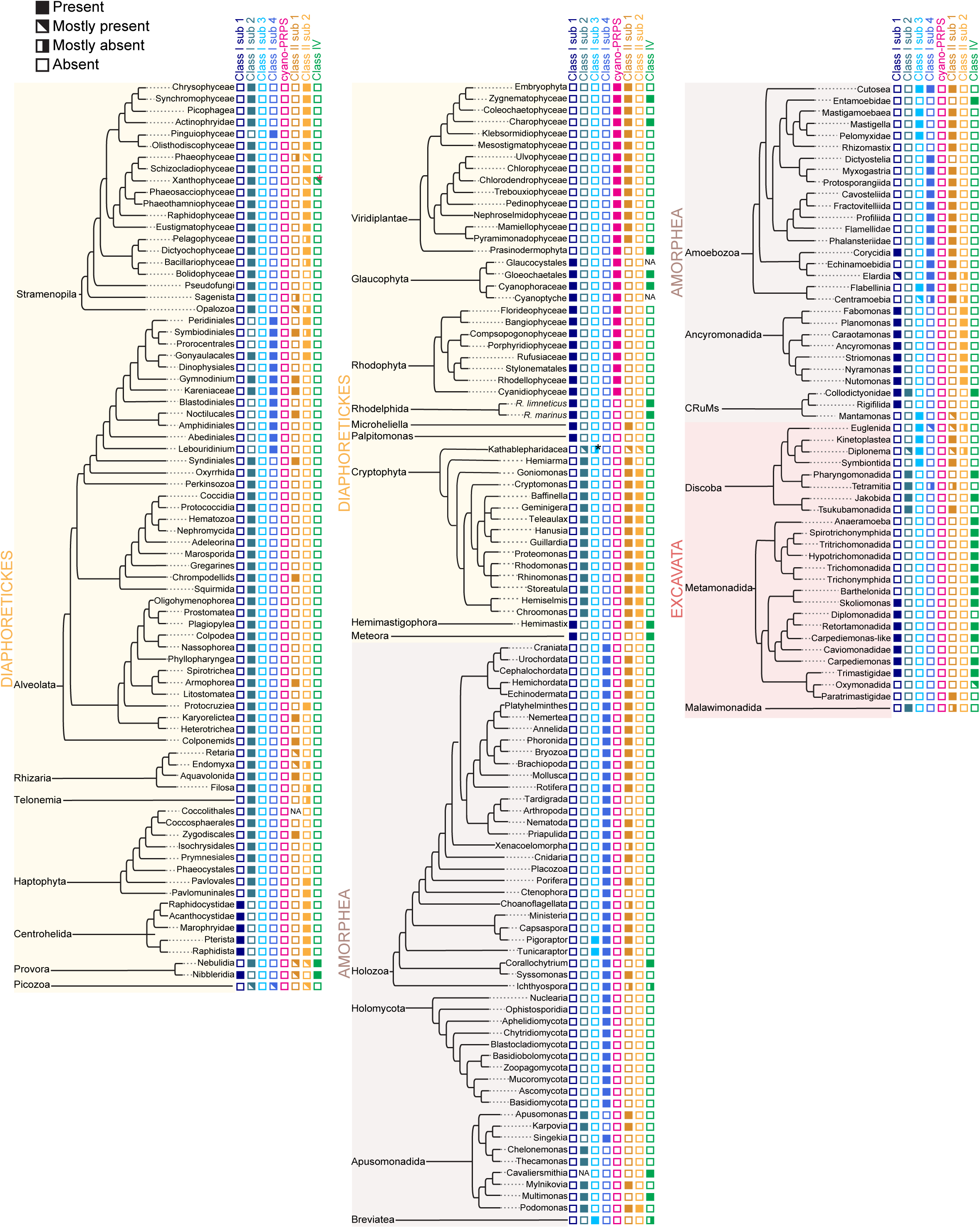
Lineage-specific PRPS gene losses drive heterogeneous distribution of PRPS classes/subclasses across eukaryotes. Expanded eukaryotic phylogenetic tree showing distribution of PRPS classes across major and minor lineages. Presence and absence are indicated using a categorical scheme (present, mostly present, mostly absent, absent) based on the proportion of sampled taxa within each lineage. “NA” denotes lineages for which presence/absence could not be determined due to incomplete or unavailable datasets. Lineages are organized by eukaryotic supergroups – Diaphoretickes, Amorphea, and Excavata – with expanded taxonomic resolution relative to Figure 1D. Colored annotations correspond to PRPS subclasses (Class I subclasses 1-4, cyano-PRPS, Class II subclasses 1-2, and Class IV). Black asterisk denotes Class II PRPS restricted to a single Cryptophyta species (uncultured Katablepharidaceae), consistent with horizontal gene transfer, and red asterisk indicates Class IV PRPS in a limited number of Stramenopila (three Xanthophyceae species), also suggestive of horizontal gene transfer.

**Supplementary Figure 6.**
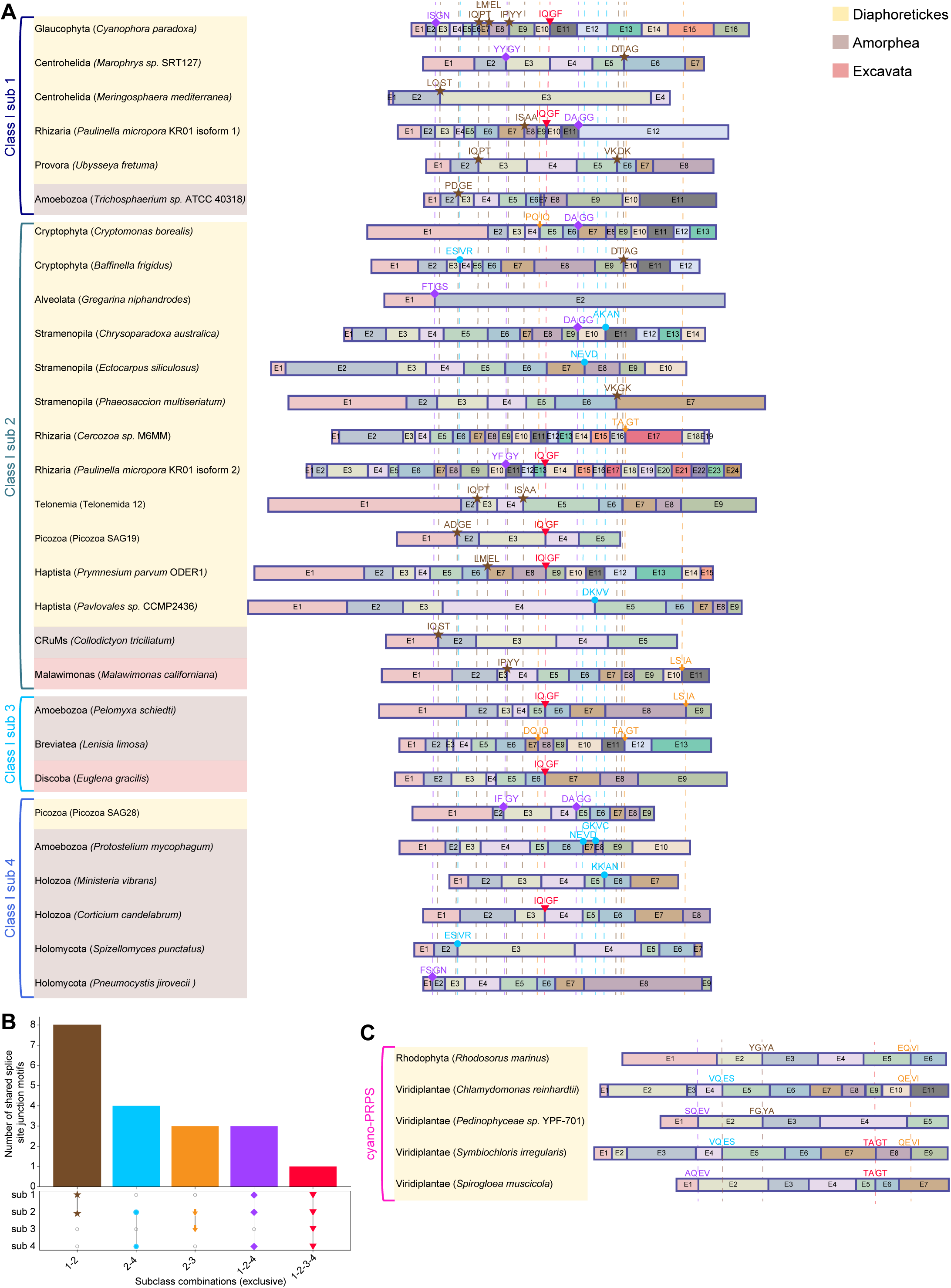
Conserved splice junctions reveal shared ancestry of Class I PRPS genes and of cyanobacteria-derived PRPS genes. **(A)** Exon organization of Class I PRPS subclasses (subclasses 1-4) across representative eukaryotic lineages, shown as colored boxes with lengths proportional to the number of amino acid residues per exon; introns are not displayed. Sequences are grouped by supergroups – Diaphoretickes, Amorphea, and Excavata. Conserved splice site junctions revealed via multiple sequence alignments of translated sequences are indicated at exon boundaries, with amino acids flanking each junction (two residues on either side) displayed. Symbols denote shared splice site junction categorized by subclass combinations as defined in (B). **(B)** Number of shared splice site junctions across Class I subclasses. Bars represent counts of conserved junctions, with subclass combinations indicated below. **(C)** Exon organization of cyano-PRPS across representative Archaeplastida species. Conserved splice site junctions revealed via multiple sequence alignments of translated sequences are indicated at exon boundaries, with amino acids flanking each junction (two residues on either side) displayed.

**Supplementary Figure 7.**
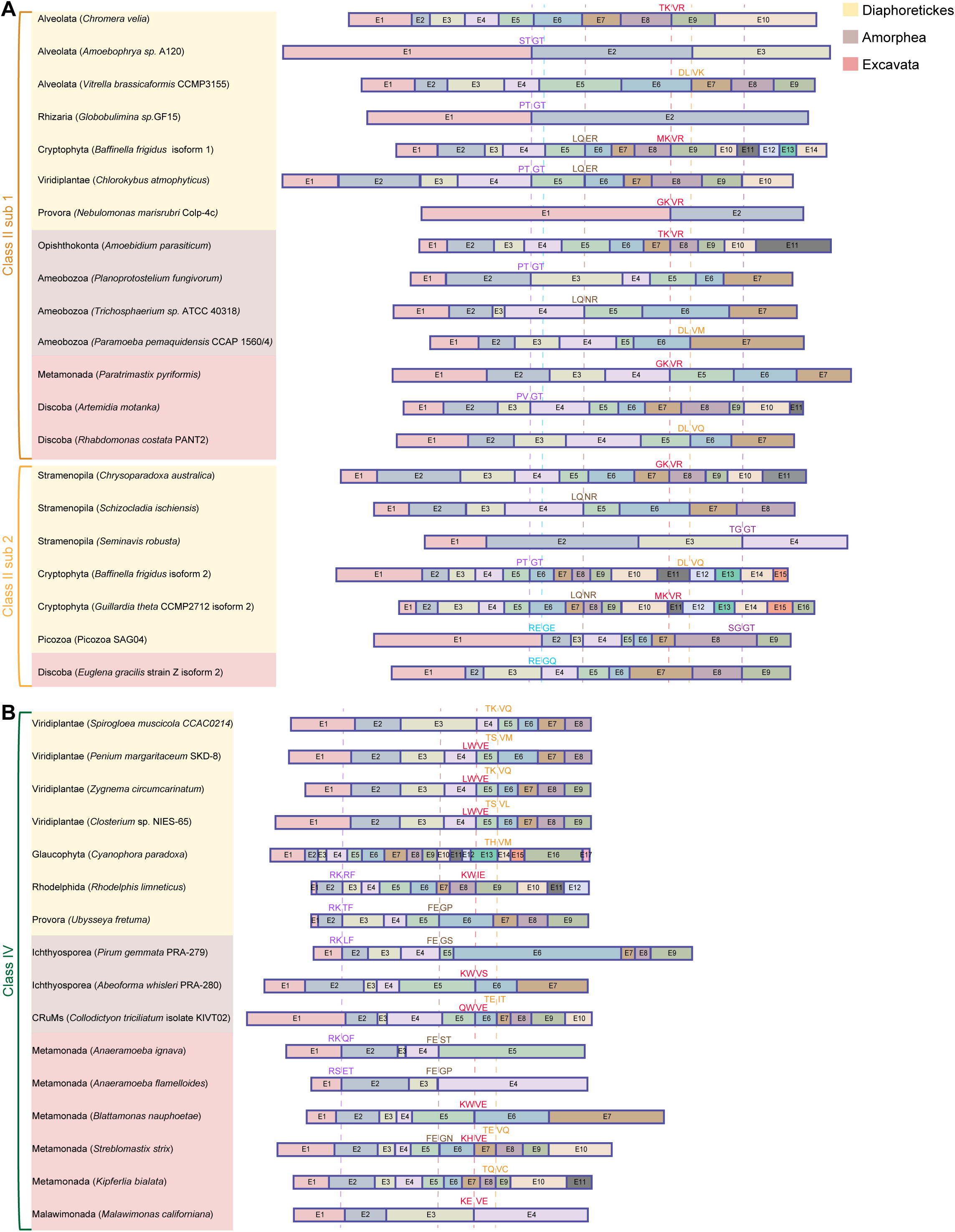
Conserved splice junctions reveal shared ancestry of Class II PRPS genes and of Class IV PRPS genes. **(A-B)** Exon organization of Class II PRPS subclasses (subclasses 1-2) (A) and Class IV PRPS (B) across representative eukaryotic lineages, shown as colored boxes with lengths proportional to the number of amino acid residues per exon; introns are not displayed. Sequences are grouped by major supergroups (Diaphoretickes, Amorphea, and Excavata). Conserved splice site junctions revealed via multiple sequence alignments of translated sequences are indicated at exon boundaries, with amino acids flanking each junction (two residues on either side) displayed.

**Supplementary Figure 8.**
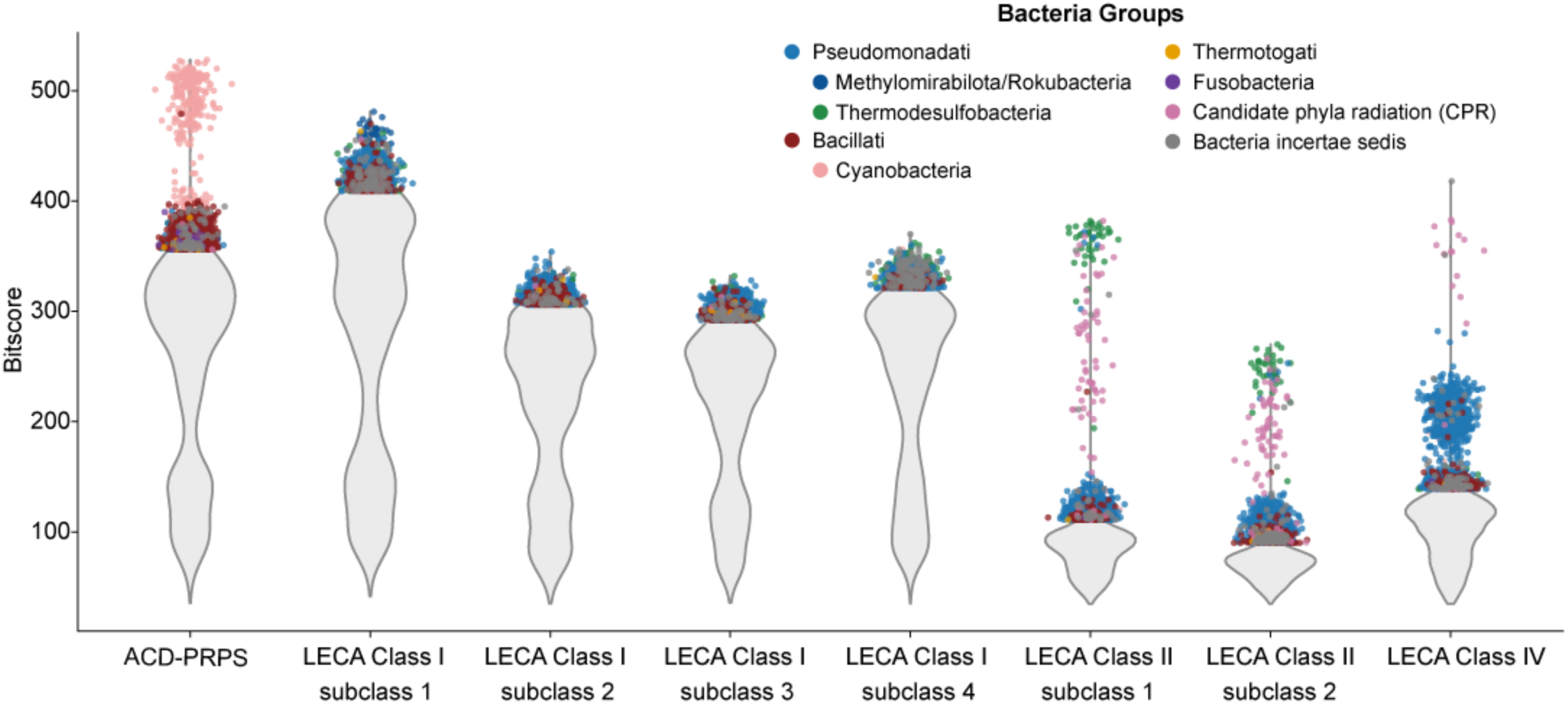
BLASTP profiling of ancestral PRPS reconstructions support chimeric bacterial origins of the pre-LECA eukaryotic PRPS repertoire. Violin plots showing the distribution of BLASTP bit scores between ancestrally reconstructed PRPS queries and a curated database of 33,248 non-redundant prokaryotic PRPS sequences (28,560 bacterial and 4,688 archaeal). Queries include ACD-PRPS, LECA Class I subclasses (subclasses 1-4), LECA Class II subclasses (subclasses 1-2), and LECA Class IV. Ancestral sequence reconstruction (ASR) is described in Methods. For each query, the top 5% of highest-scoring sequences (≥ 95th percentile) are overlaid as individual points and colored by taxonomic classification (Pseudomonadati, Methylomirabilota/Rokubacteria, Bacillati, Cyanobacteria, Thermotogati, Fusobacteria, Candidate Phyla Radiation (CPR), and Bacteria incertae sedis).

**Supplementary Figure 9.**
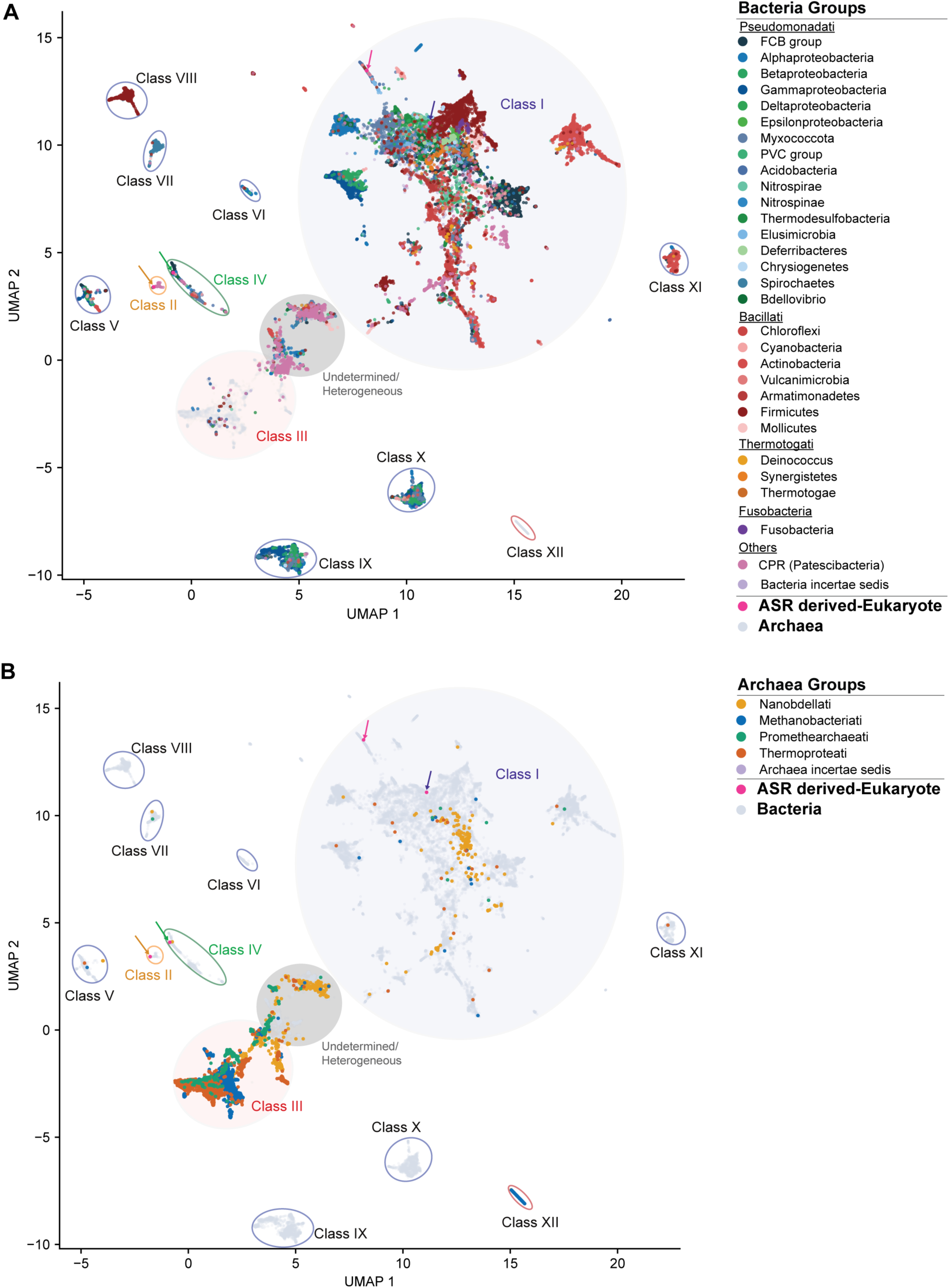
Class I and Class III PRPS have near universal distribution in Bacteria and Archaea, respectively, whereas other classes are more sporadically distributed. **(A-B)** Two-dimensional UMAP of ESM-2-derived embeddings for prokaryotic PRPS sequences (same embedding as Figure 2A). Sequences are colored by kingdom/phylum-level groupings for bacterial lineage in (A) and archaeal lineage in (B), respectively, indicating the relative distribution of sequences across major bacterial and archaeal groups. Different prokaryotic PRPS classes are indicated and circled. Colored arrows denote ancestrally reconstructed eukaryotic sequences (blue, LECA Class I sub 1; pink, ACD-PRPS; orange, LECA Class II sub 1; green, LECA Class IV).

**Supplementary Figure 10.**
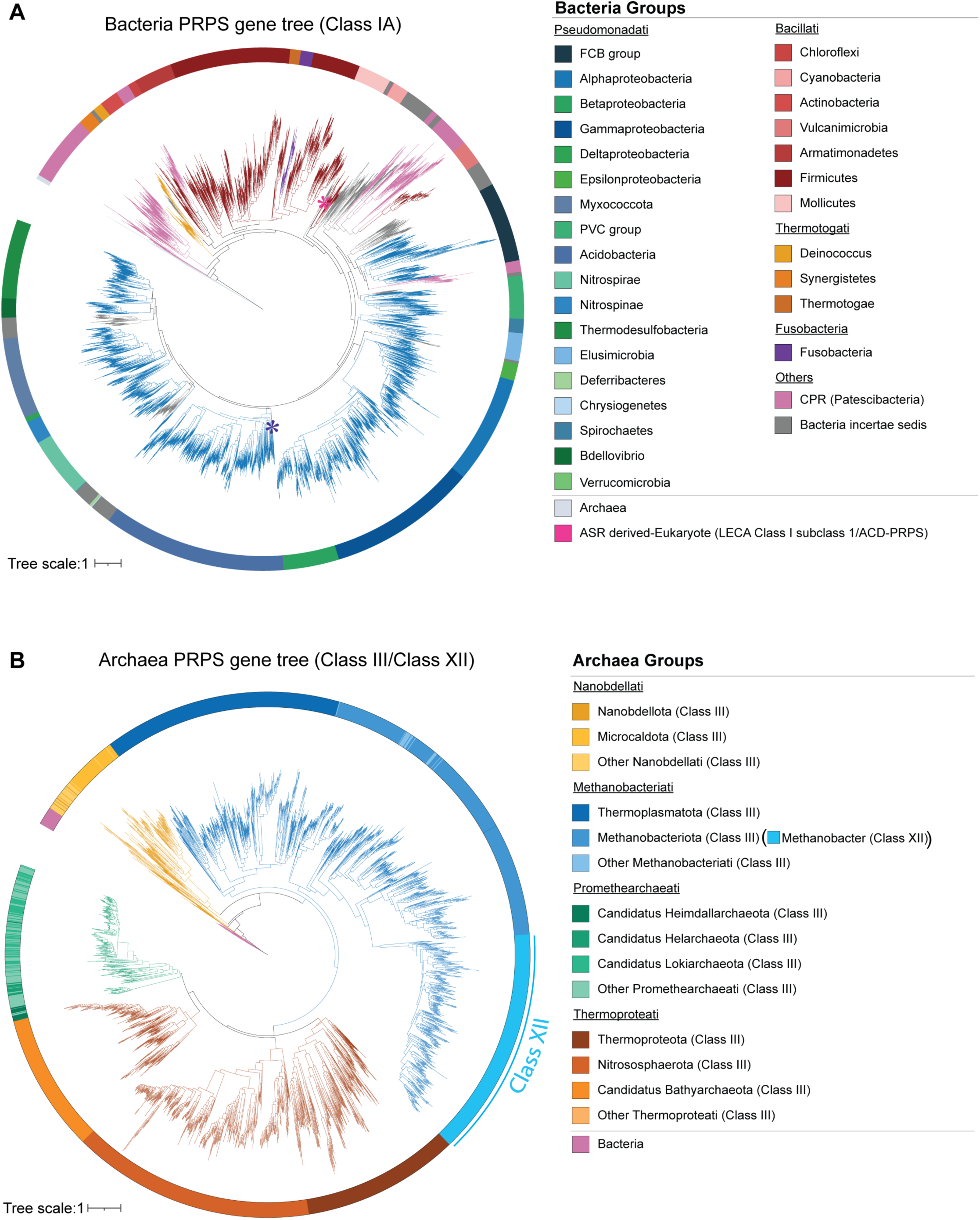
Phylogenetics indicate vertical inheritance of the most prevalent bacterial (Class I) and archaeal (Class III) PRPS classes. **(A)** Circular maximum likelihood gene tree of bacterial Class I PRPS sequences, inferred using IQ-TREE under the LG+F+R10 model. Branches are colored by major bacterial lineages, and the outer circular color strip denotes kingdom/phylum-level classification. Archaeal sequences are included as an outgroup. The most prevalent Class I PRPS sequences that follow a pattern of vertical inheritance within bacteria are hereafter referred to as Class IA. **(B)** Circular maximum likelihood gene tree of archaeal PRPS sequences, including Class III and Class XII classes, inferred using IQ-TREE under the LG+F+R10 model. Branches are colored by major archaeal lineages, and the outer circular color strip denotes kingdom/phylum-level classification. Class XII sequences are labeled in the circular color strip. Bacterial Class IA sequences are included as an outgroup. The full tree for (A) and (B) is provided in rectangular format with bootstrap support values in Supplementary File 2 and 3, respectively, and in Newick format in the Figshare repository.

**Supplementary Figure 11.**
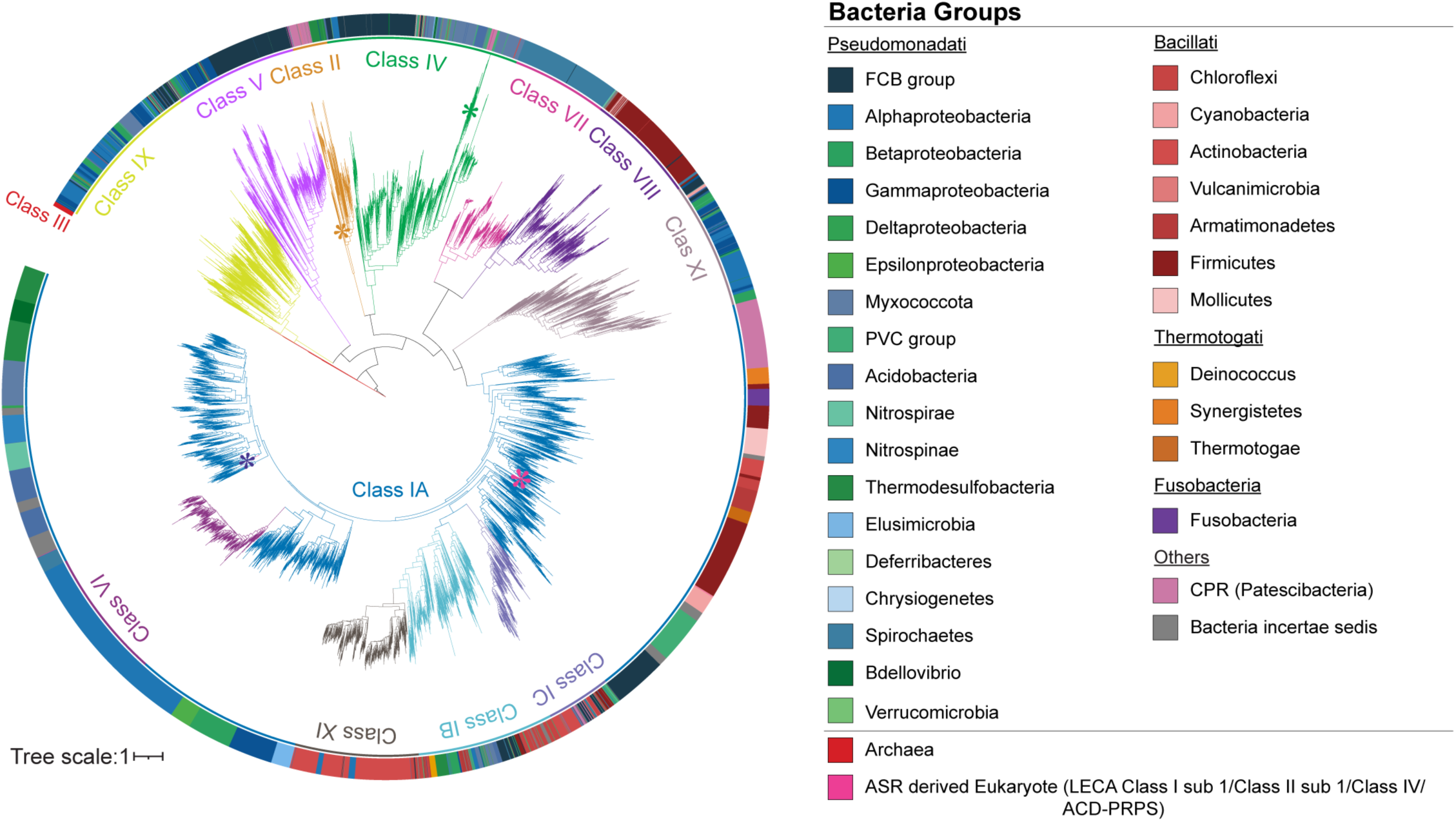
Expanded gene tree reveals relationships among prokaryotic PRPS classes. Circular maximum likelihood gene tree of diverse bacterial (Class I (A, B, C), II, IV-XI) and archaeal (Class III) PRPS sequences inferred using IQ-TREE under the LG+F+R10 model (same as Figure 2C). Branches are colored based on PRPS classes as indicated. Outer circular color strip denotes kingdom/phylum-level classification. Asterisks mark positions of ASR-derived ancestral sequences (blue, LECA Class I sub 1; pink, ACD-PRPS; green, LECA Class IV; orange, LECA Class II sub 1). The full tree is provided in rectangular format with bootstrap support values in Supplementary File 5, and in Newick format in the Figshare repository.

**Supplementary Figure 12.**
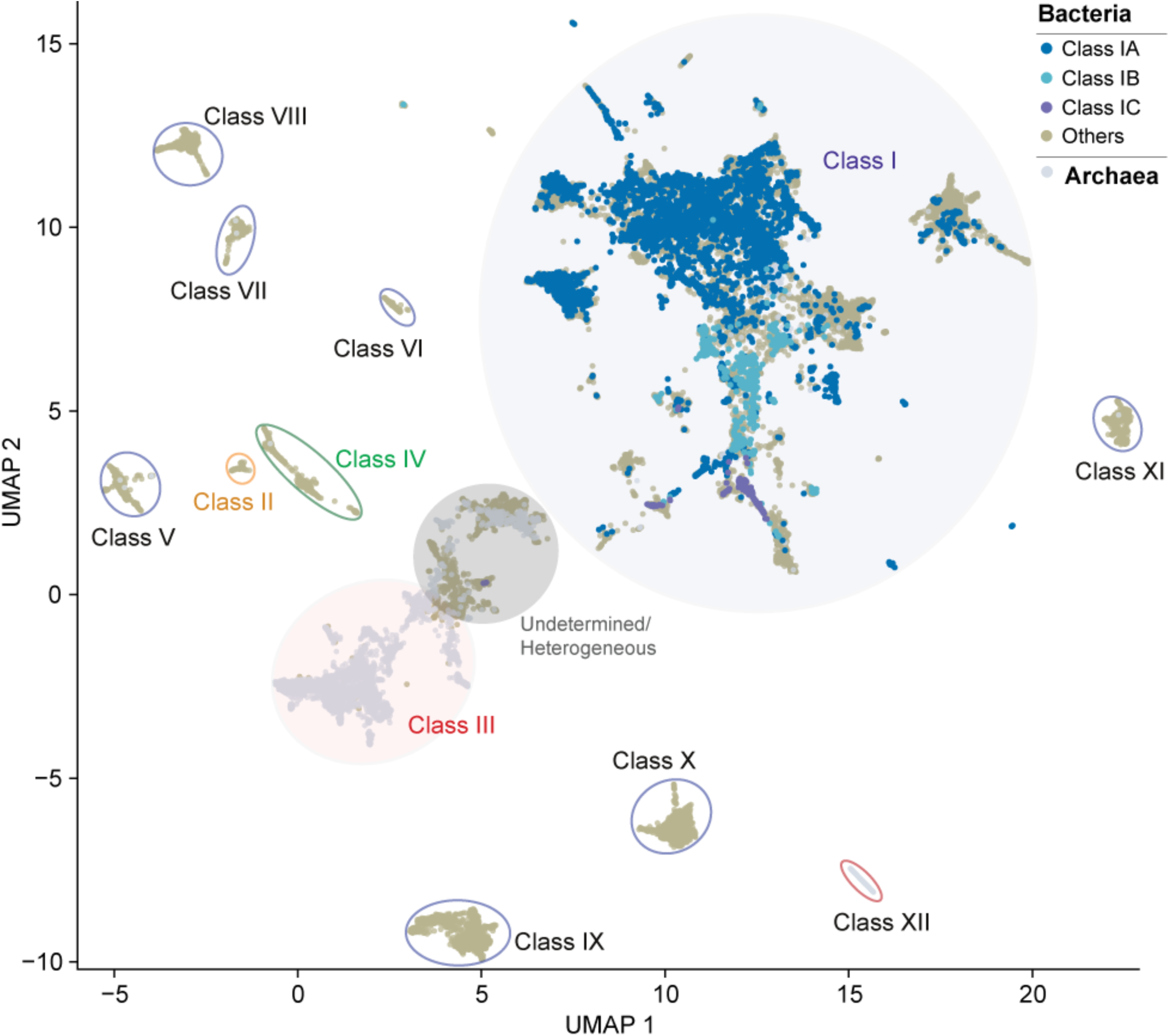
Bacterial Class I subclasses occupy distinct regions within the broader Class I embedding landscape. Two-dimensional UMAP of ESM-2-derived embeddings for prokaryotic PRPS sequences (same embedding as Figure 2A) highlighting bacterial Class I subclass – Class IA (n = 5,224), IB (n = 699), and IC (n = 174). Different prokaryotic PRPS classes are indicated and circled.

**Supplementary Figure 13.**
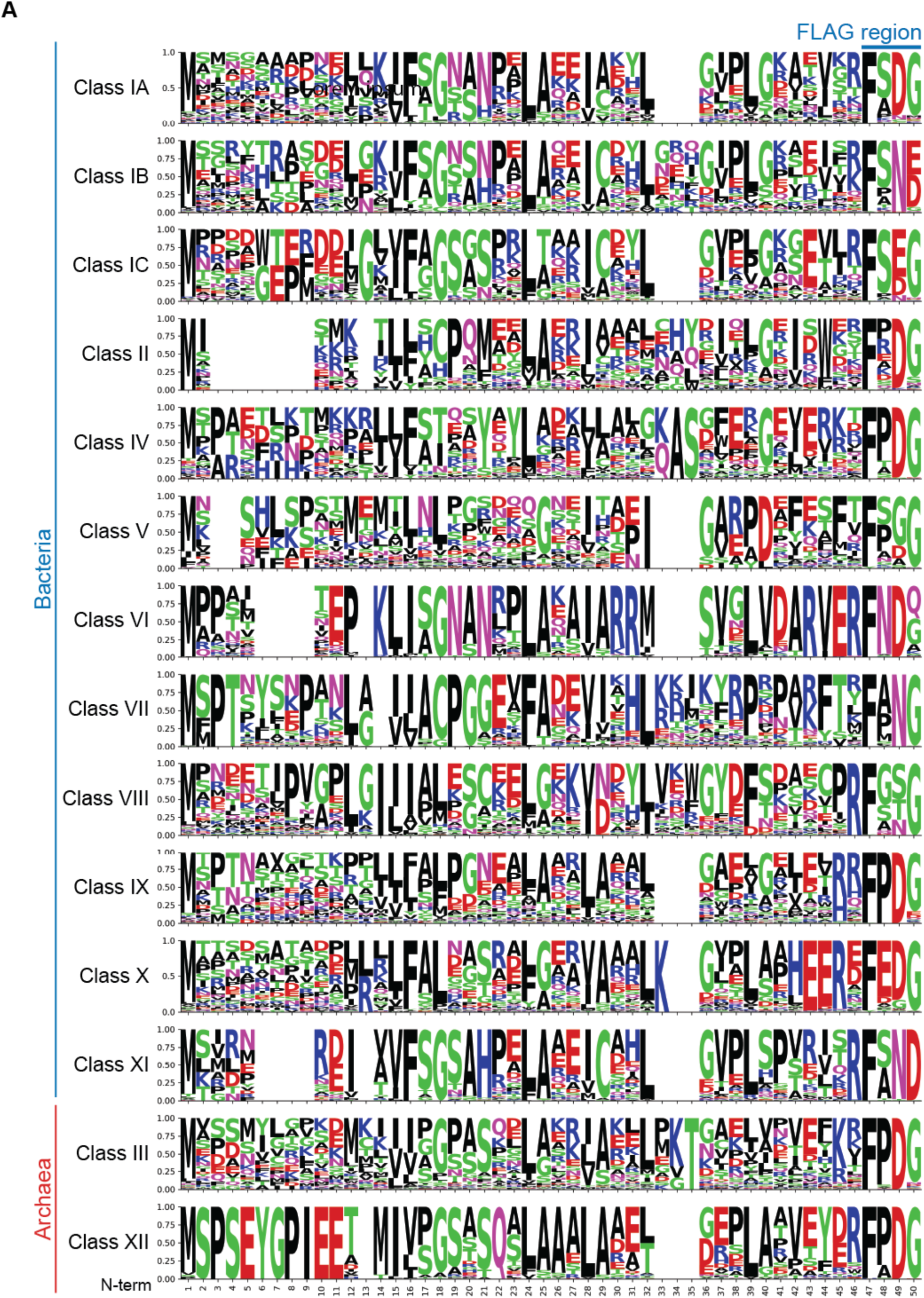
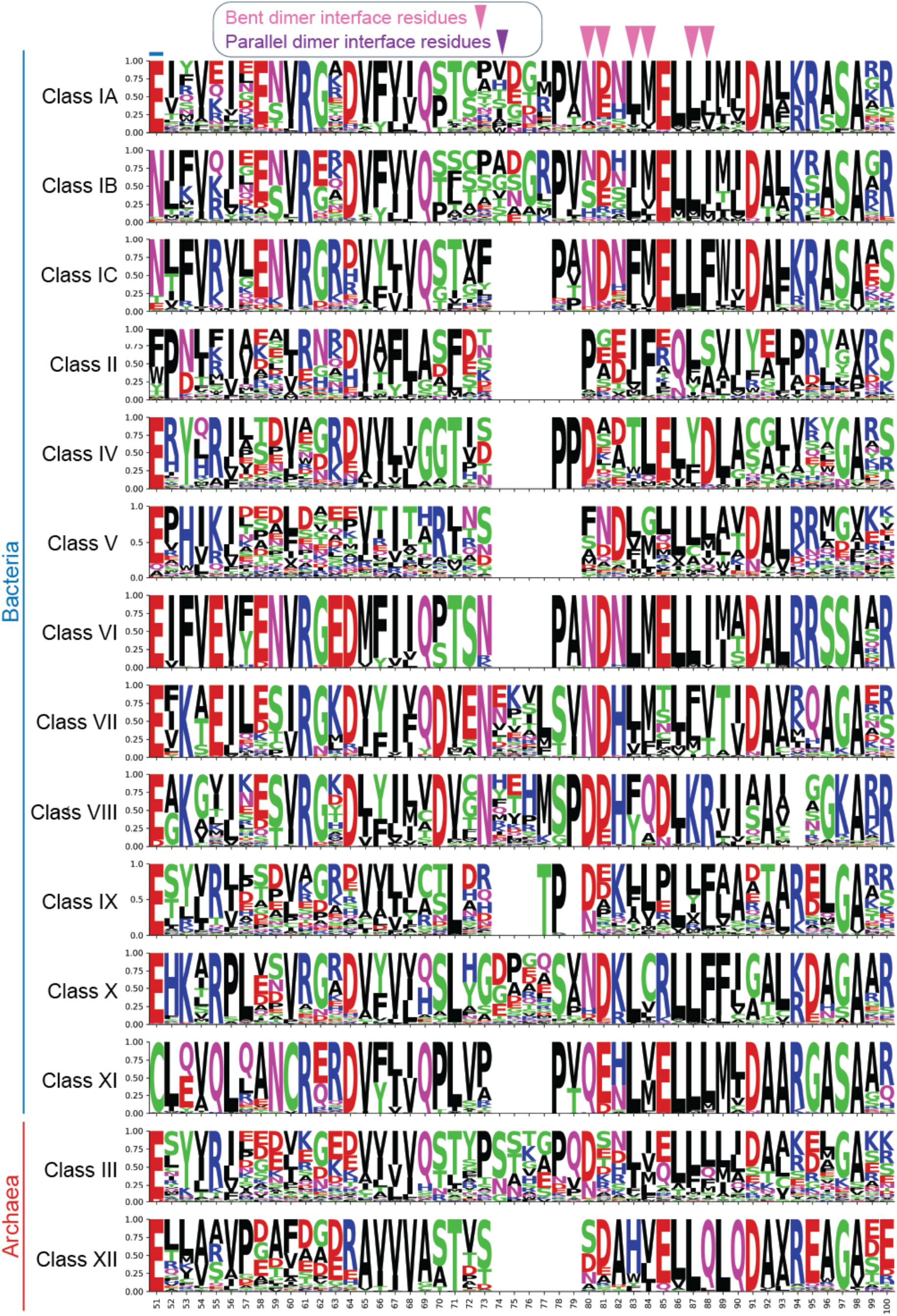
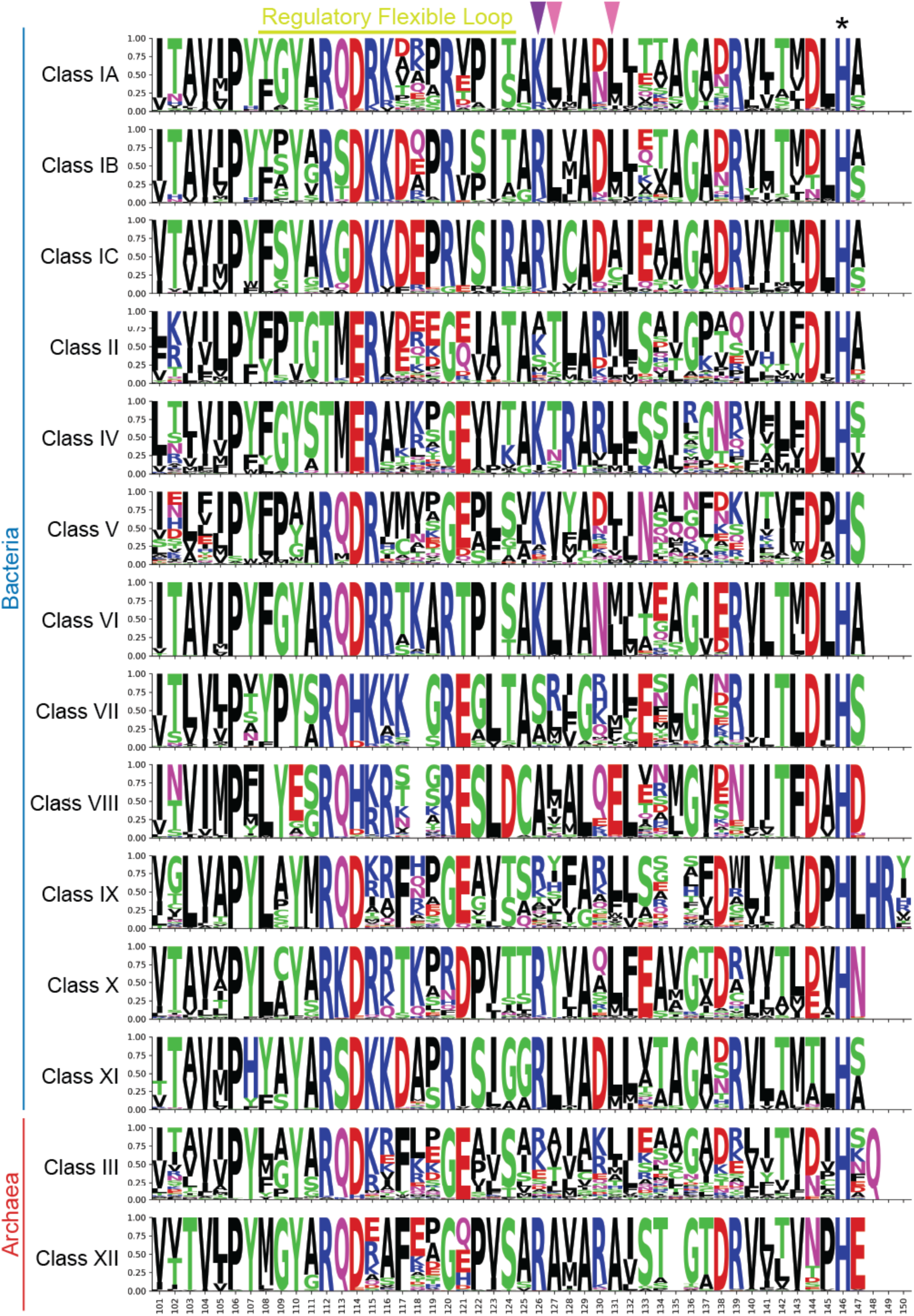
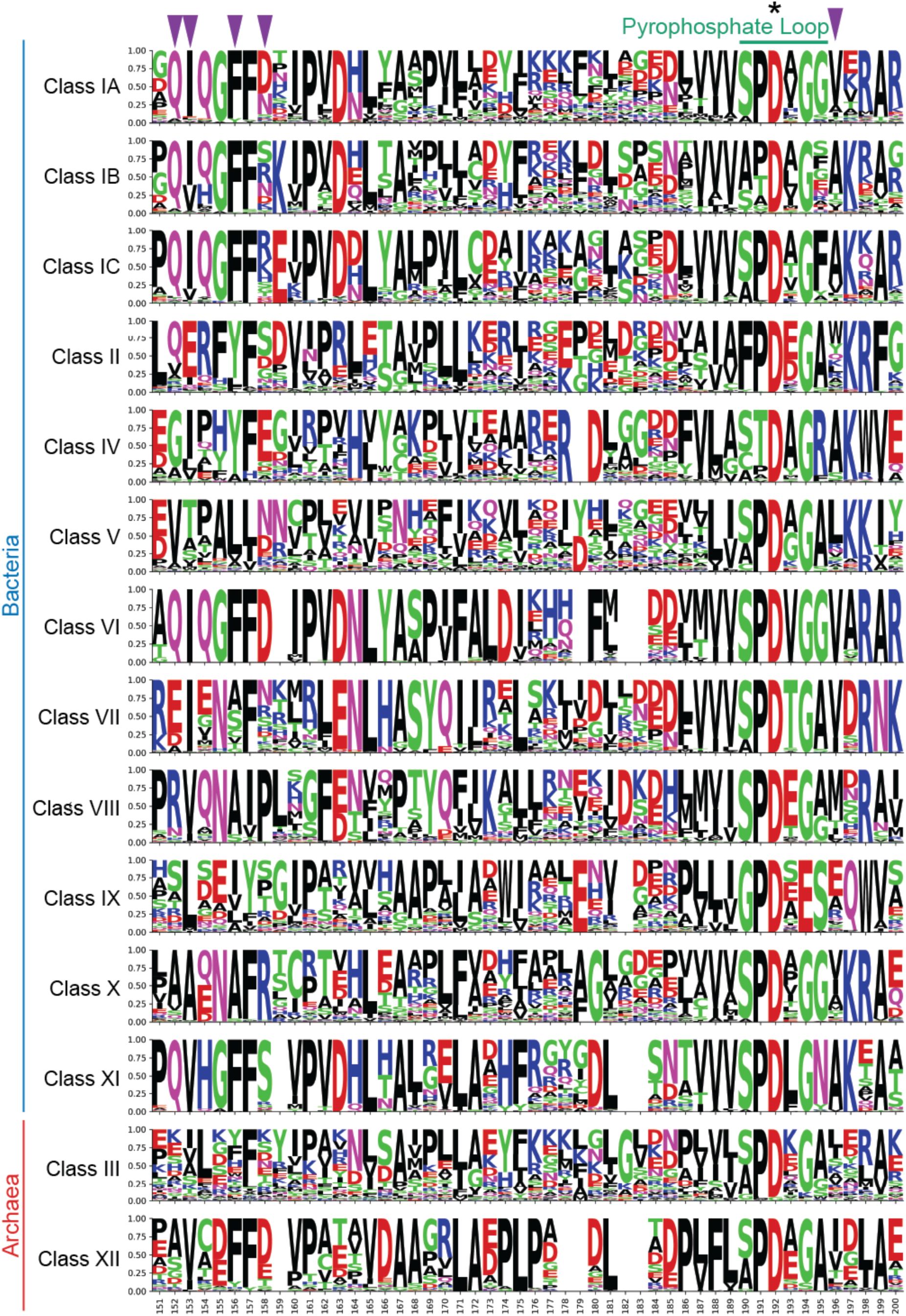
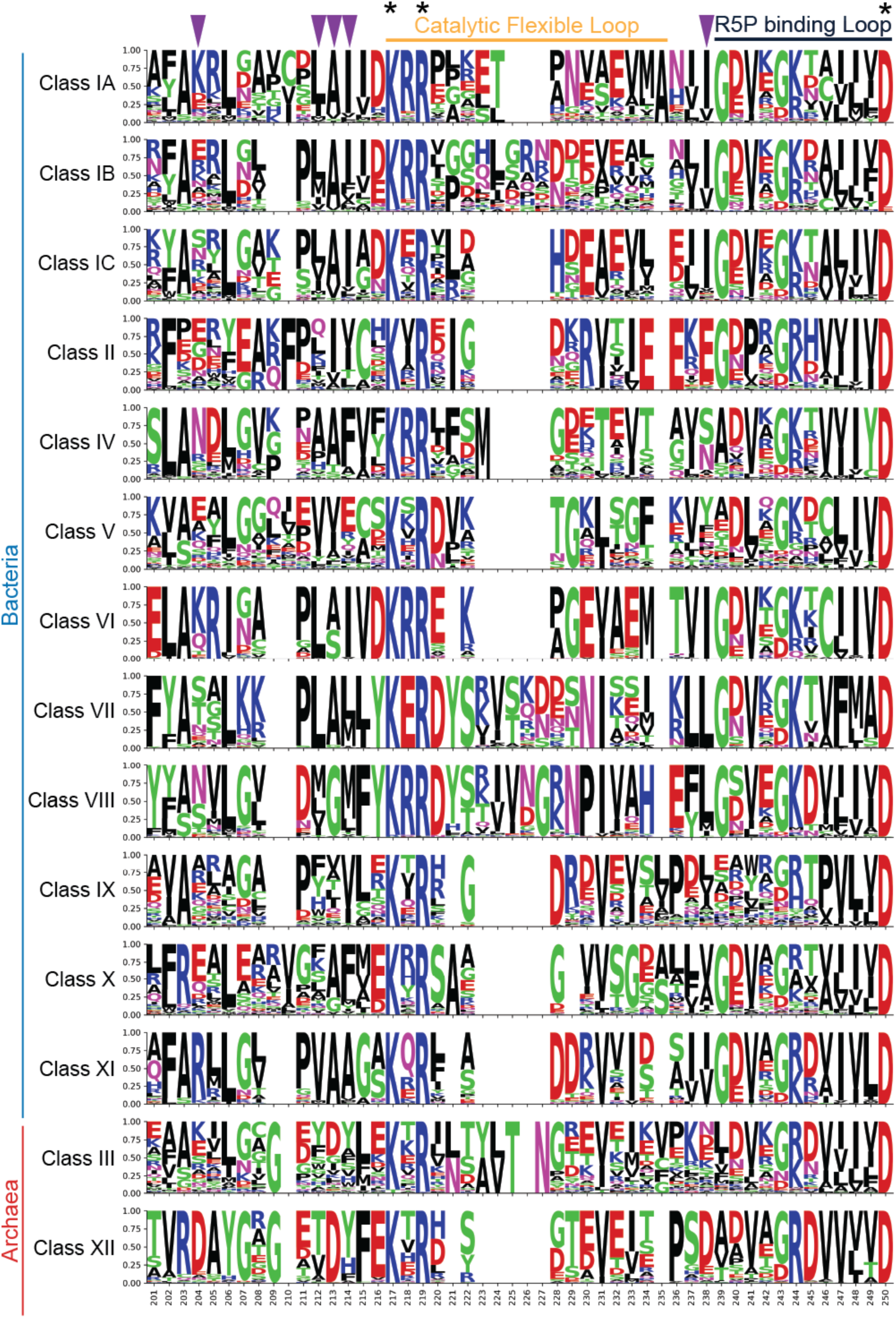
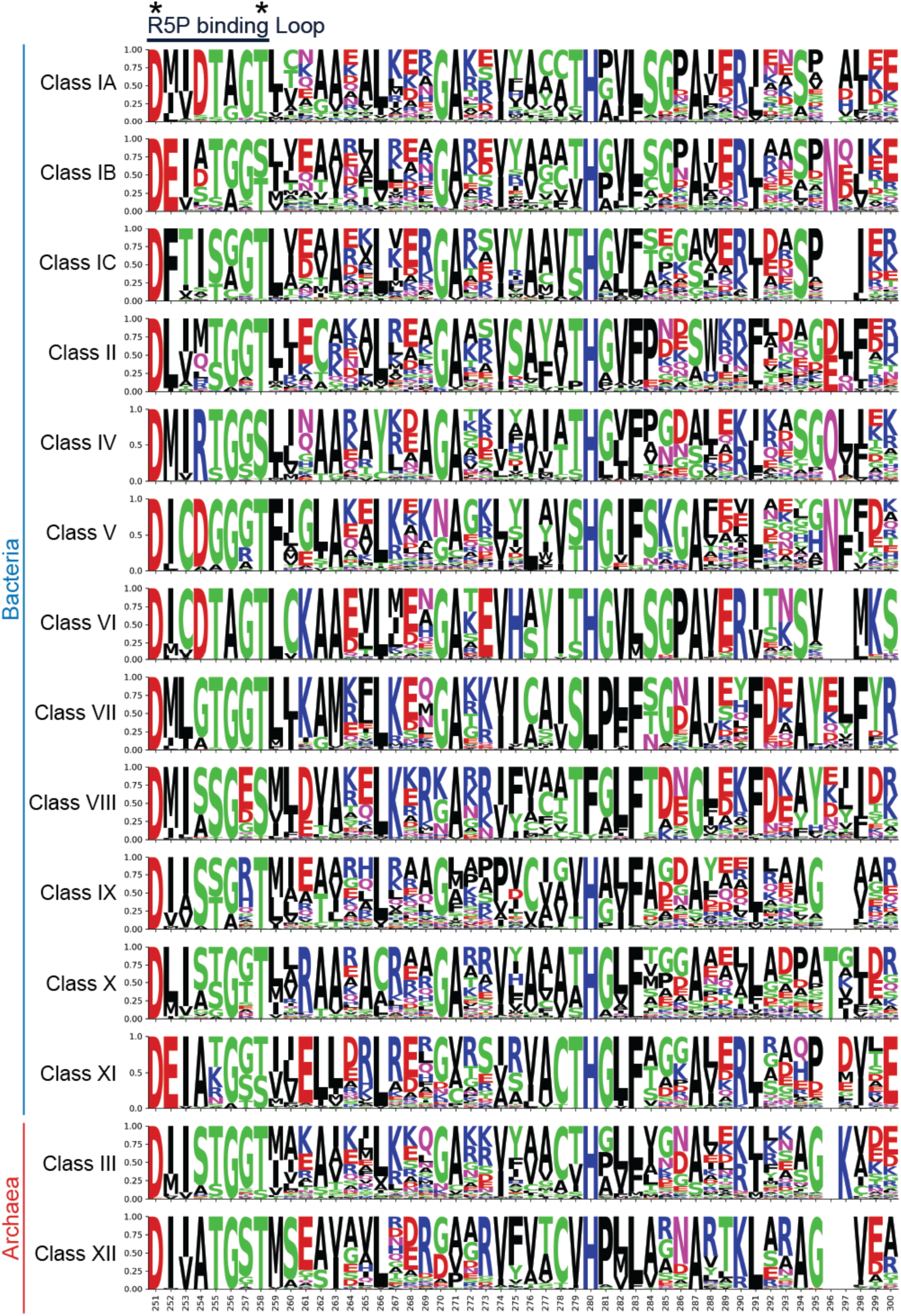
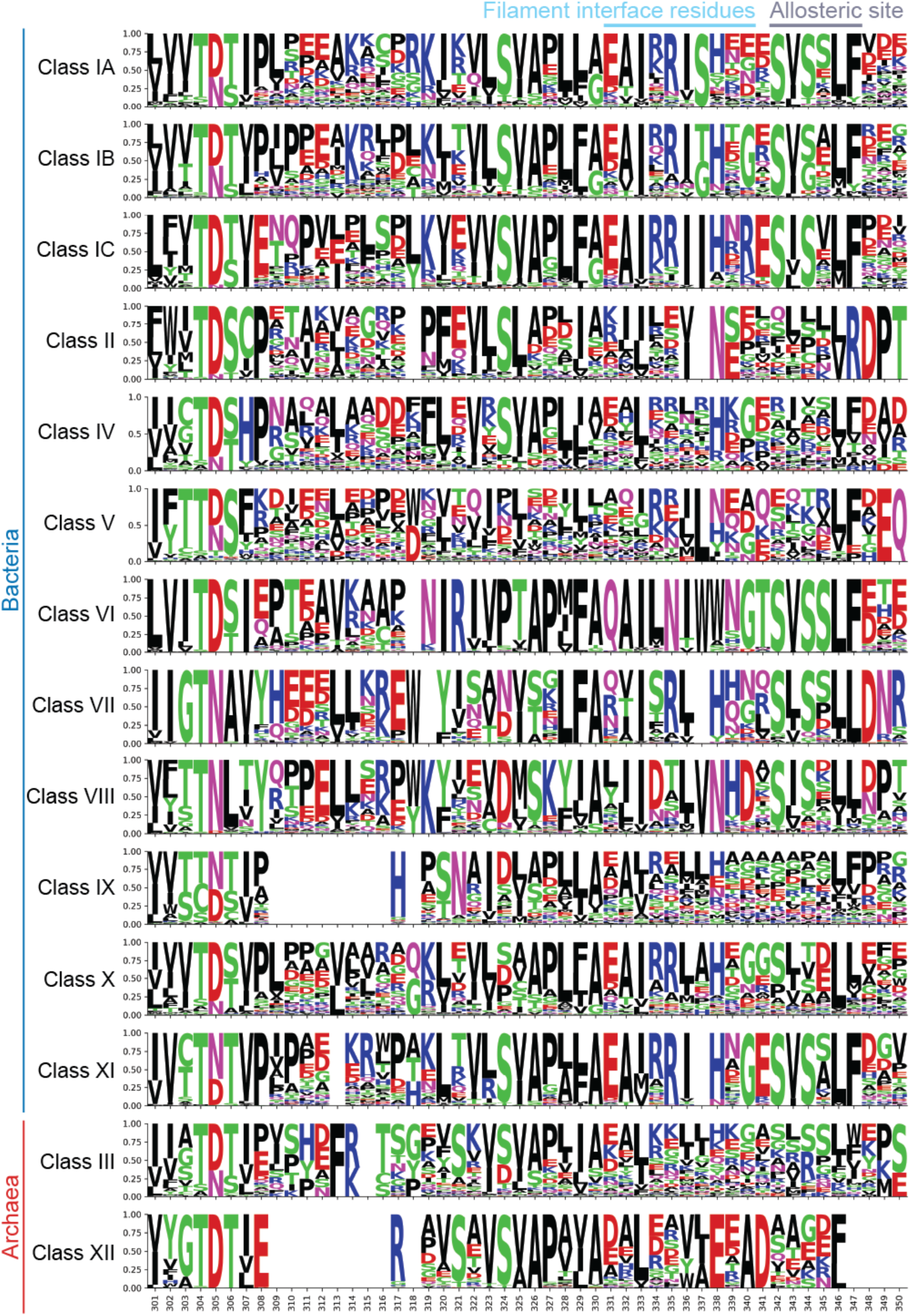
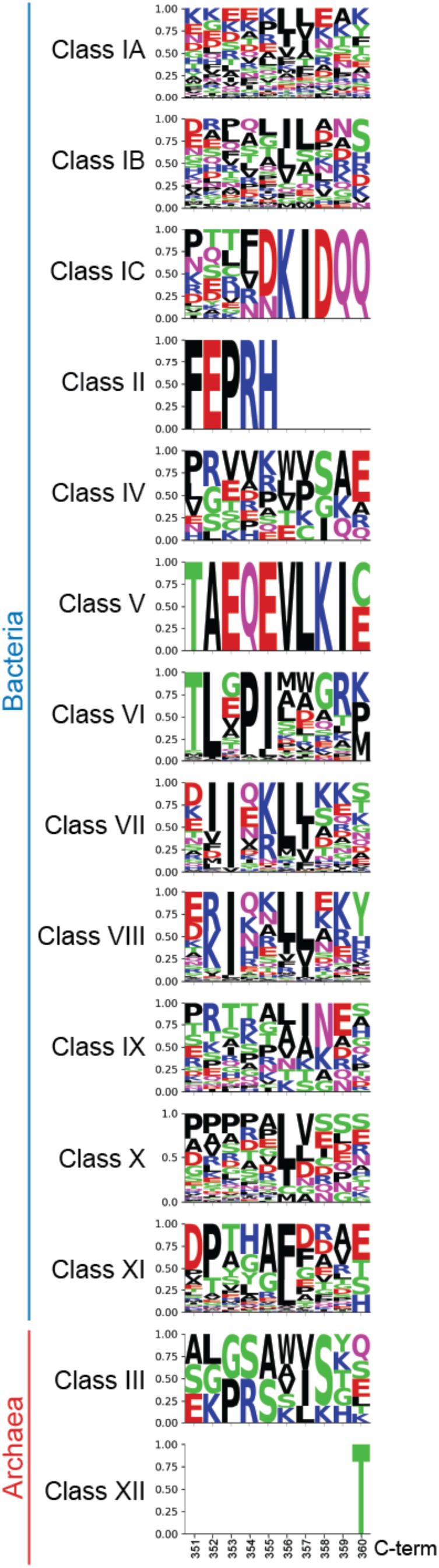
Global sequence conservation and divergence across prokaryotic PRPS classes. WebLogo depicting full-length multiple sequence alignment for prokaryotic PRPS sequences. Logos represent amino acid frequencies at each alignment position, calculated independently for each group using non-gap residues. The height of each letter reflects its relative frequency at that position. Asterisks denote universally conserved residues critical for enzyme function. Group sizes for bacterial PRPS classes: Class IA (n = 3251), Class IB (n = 732), Class IC (n = 224), Class II (n = 109), Class IV (n = 763), Class V (n = 511), Class VI (n = 459), Class VII (n = 445), Class VIII (n = 726), Class IX (n = 1368), Class X (n = 929), Class XI (n = 699). Group sizes for archaeal PRPS classes: Class III (n = 1156), Class XII (n = 410).

**Supplementary Figure 14.**
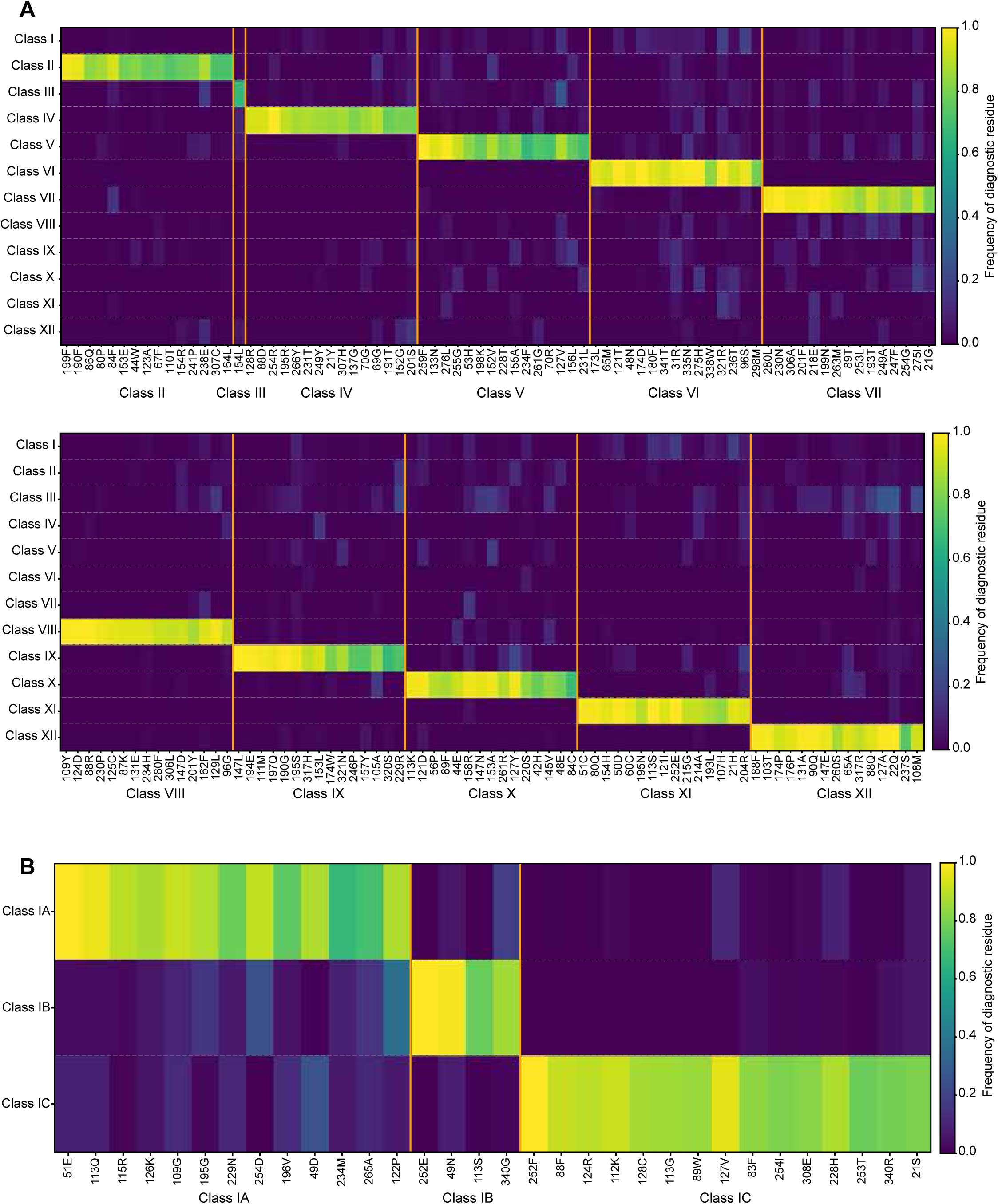
Diagnostic residue signatures define distinct PRPS classes across prokaryotes. **(A-B)** Heatmaps showing diagnostic residues distinguishing PRPS classes across Bacteria and Archaea (A) and across bacterial Class I subclasses (B). Rows correspond to PRPS classes and columns represent alignment positions meeting stringent diagnostic criteria (frequency, coverage, and statistical criteria – see Methods). Cells indicate residue frequency (0-1) within each group, calculated from non-gap residues at each position. Only the top diagnostic sites per class are shown. Diagnostic site positions for each class are referenced to amino acid position shown in Supplementary Figure 13. Group sizes for (A): Class I (n = 4171), Class II (n = 109), Class III (n = 1156), Class IV (n = 763), Class V (n = 511), Class VI (n = 459), Class VII (n = 445), Class VIII (n = 726), Class IX (n = 1368), Class X (n = 929), Class XI (n = 699), Class XII (n = 410). Group sizes for (B): Class IA (n = 3251), Class IB (n = 732), Class IC (n = 224).

**Supplementary Figure 15.**
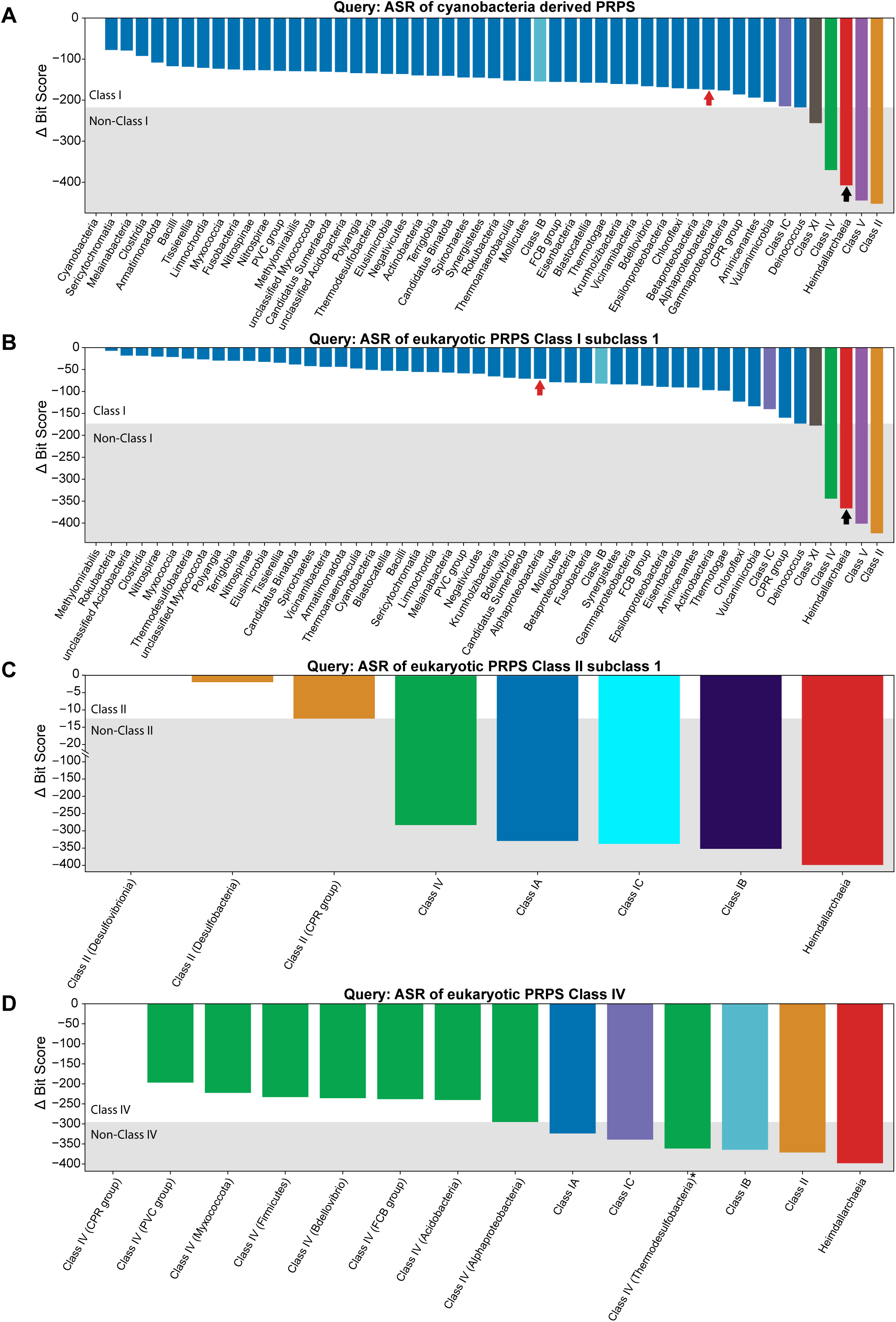
Ancestrally reconstructed eukaryotic PRPS sequences nominate distinct bacterial lineages as HGT donors via HMM profiling. **(A-D)** Waterfall plots showing Δ bit score profiles for ancestrally reconstructed ACD-PRPS (A), LECA Class I subclass 1 (B), LECA Class II subclass 1 (C), and LECA Class IV (D) sequences against a panel of lineage-specific profile Hidden Markov Models (HMMs). For each query, scores were normalized to the top-scoring model (Δ = 0), and bars are ordered by decreasing bit score. The shaded region denotes non-Class I PRPS. In (A) and (B), ACD-PRPS and LECA Class I show highest-scoring matches to profile HMMs of Cyanobacteria and Methylomirabilota respectively. Alphaproteobacterial HMM (red arrows) and archaeal HMM (black arrows) are not among the top-scoring profiles suggesting that they are unlikely to be the donors to stem eukaryote. In (C) and (D), LECA Class II and LECA Class IV show highest-scoring matches to profile HMMs of Thermodesulfobacteria and Candidate Phyla Radiation (CPR), respectively. Archaea ranks lowest among the profiles compared. Black asterisk denotes a divergent Class IV sequence in Thermodesulfobacteria. Ancestral sequence reconstruction (ASR) is described in Methods.

**Supplementary Figure 16.**
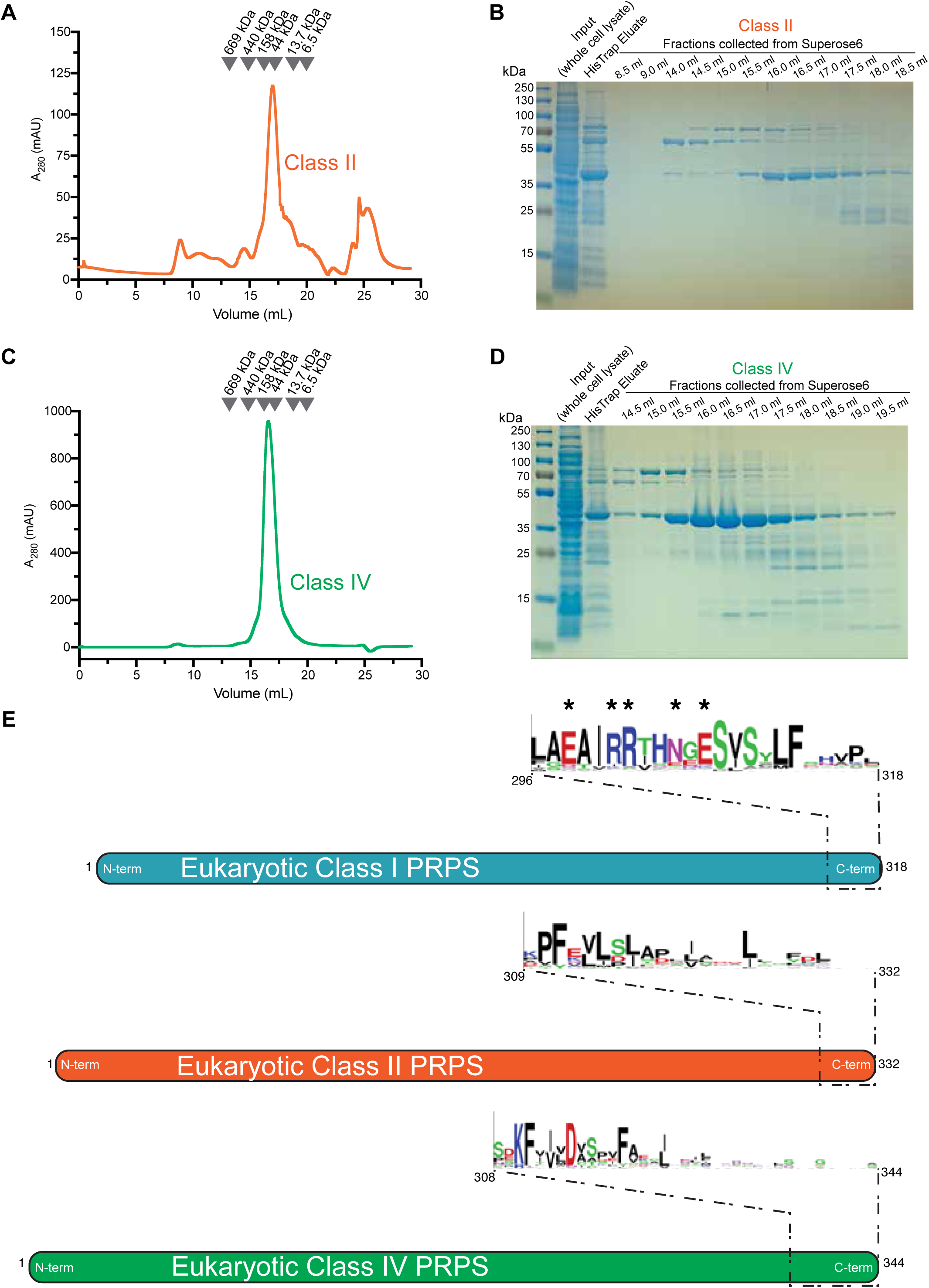
Purification and determination of oligomeric state of recombinant Class II and Class IV PRPS. **(A)** Size-exclusion chromatography (SEC) elution profile of recombinant protein purified from bacteria for *Branchiostoma lanceolatum* Class II PRPS on a Superose 6 Increase 10/300 GL column; molecular weight standards are shown. **(B)** SDS-PAGE followed by Coomassie staining of fractions collected from the SEC run in (A), showing enrichment and purity of Class II PRPS in peak fractions. **(C)** SEC elution profile of recombinant protein purified from bacteria for *Corallochytrium limacisporum* Class IV PRPS on a Superose 6 Increase 10/300 GL column; molecular weight standards are shown. **(D)** SDS- PAGE followed by Coomassie staining of fractions collected from the SEC run in (C), showing enrichment and purity of Class IV PRPS in peak fractions. **(E)** WebLogo representations of conserved residues at the C-terminal region of eukaryotic PRPS, including Class I (n = 408), Class IV (n = 76), and Class II (n = 475). Residue positions are referenced to representative sequences shown in each schematic. Asterisks denote residues implicated in hexamer stacking at the filament interface for Class I PRPS^64^. These residues are conserved in Class I PRPS but are not conserved or are absent in Class IV and Class II PRPS, consistent with loss of filament-forming capability in these classes.

**Supplementary Figure 17.**
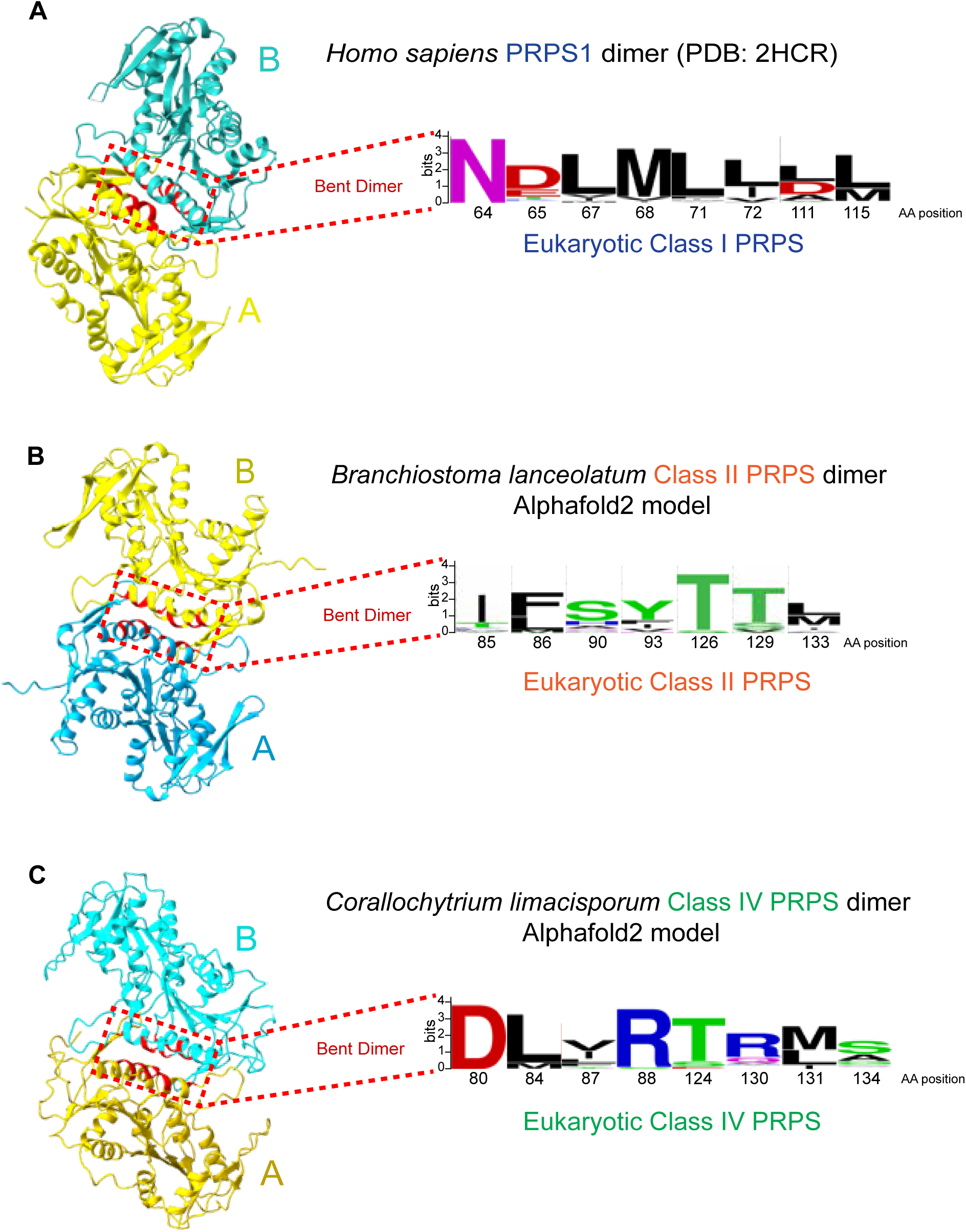
Sequence divergence at the dimer interface correspond to class-specific assembly of Class I, II, and IV PRPS. **(A)** Dimeric structure of human Class I PRPS1 (PDB: 2HCR), with the bent dimer interface highlighted (dashed box). Residues contributing to the interface between subunits A and B are shown in red. WebLogo derived from multiple sequence alignment of eukaryotic Class I PRPS sequences (n = 408) illustrates conservation of interface residues, primarily comprising helices α2 and α3, with positions referenced to *Homo sapiens* PRPS1. **(B, C)** Predicted dimeric structures of *Branchiostoma lanceolatum* Class II PRPS (B) and *Corallochytrium limacisporum* Class IV PRPS (C), generated using AlphaFold2. Interface residues from corresponding positions as shown in (A) are highlighted. WebLogo representations from eukaryotic sequences (Class IV, n = 76; Class II, n = 475) show class-specific conservation patterns within the interface region. Residue positions are referenced to the respective representative sequences.

**Supplementary Figure 18:**
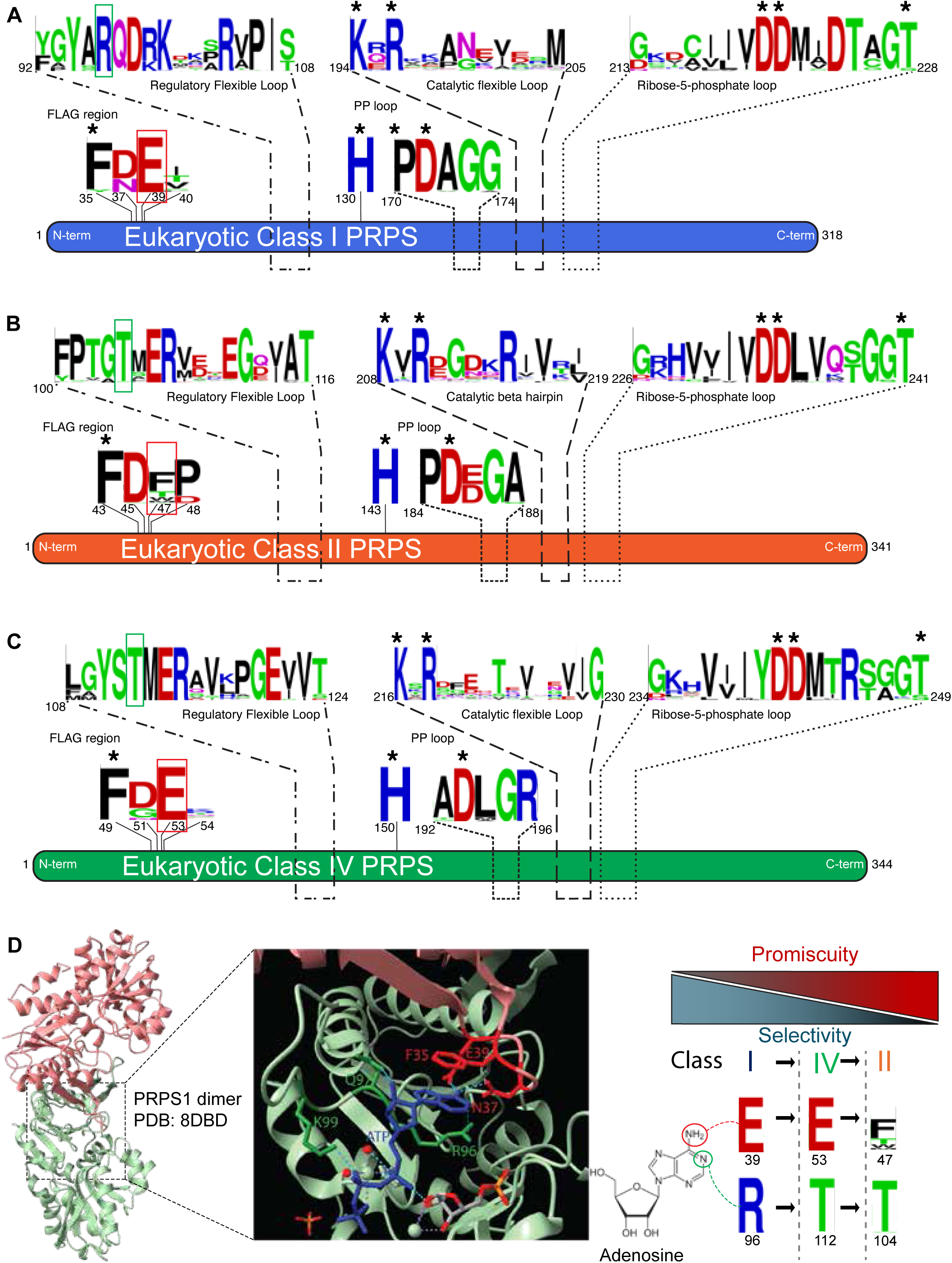
Neofunctionalization toward substrate promiscuity in the evolution of PRPS Class II. **(A-C)** representations of conserved motifs derived from multiple sequence alignments of eukaryotic PRPS, including Class I (A; n = 408), Class II (C; n = 475), and Class IV (B; n = 76). Conserved residues within key functional regions – the FLAG region, regulatory flexible (RF) loop, pyrophosphate (PP) loop, catalytic flexible (CF) loop (or catalytic β-hairpin in Class II), and ribose-5-phosphate (R5P) binding loop are indicated. Residue positions refer to the following representative sequences: *Homo sapiens* PRPS1 for Class I, *Amoebidium parasiticum* PRPS for Class II, and *Corallochytrium limacisporum* PRPS for Class IV. Asterisks denote conserved catalytic residues shared across PRPS classes. Comparative motif composition highlights divergence in regulatory and catalytic regions, with Class II exhibiting distinct sequence features consistent with altered substrate interactions. Green and red boxes highlight key residues in the RF loop and FLAG region that confer ATP specificity. **(D)** Structural basis of catalytic site divergence. A PRPS1 dimer (PDB: 8DBD) is shown (left) with monomers colored in light green and salmon. A zoomed-in view of the active site highlights interactions of ATP with FLAG and RF loop residues, with key side chains labeled (middle). Structure of adenosine is shown to emphasize specific interactions with key Class I residues and a stepwise shift away from ATP specificity (right). The gradient illustrates a conceptual shift from a specialist enzyme specific to ATP as a diphosphoryl donor to a generalist enzyme with increased substrate promiscuity (Class II), with Class IV exhibiting intermediate characteristics.

**Supplementary Figure 19.**
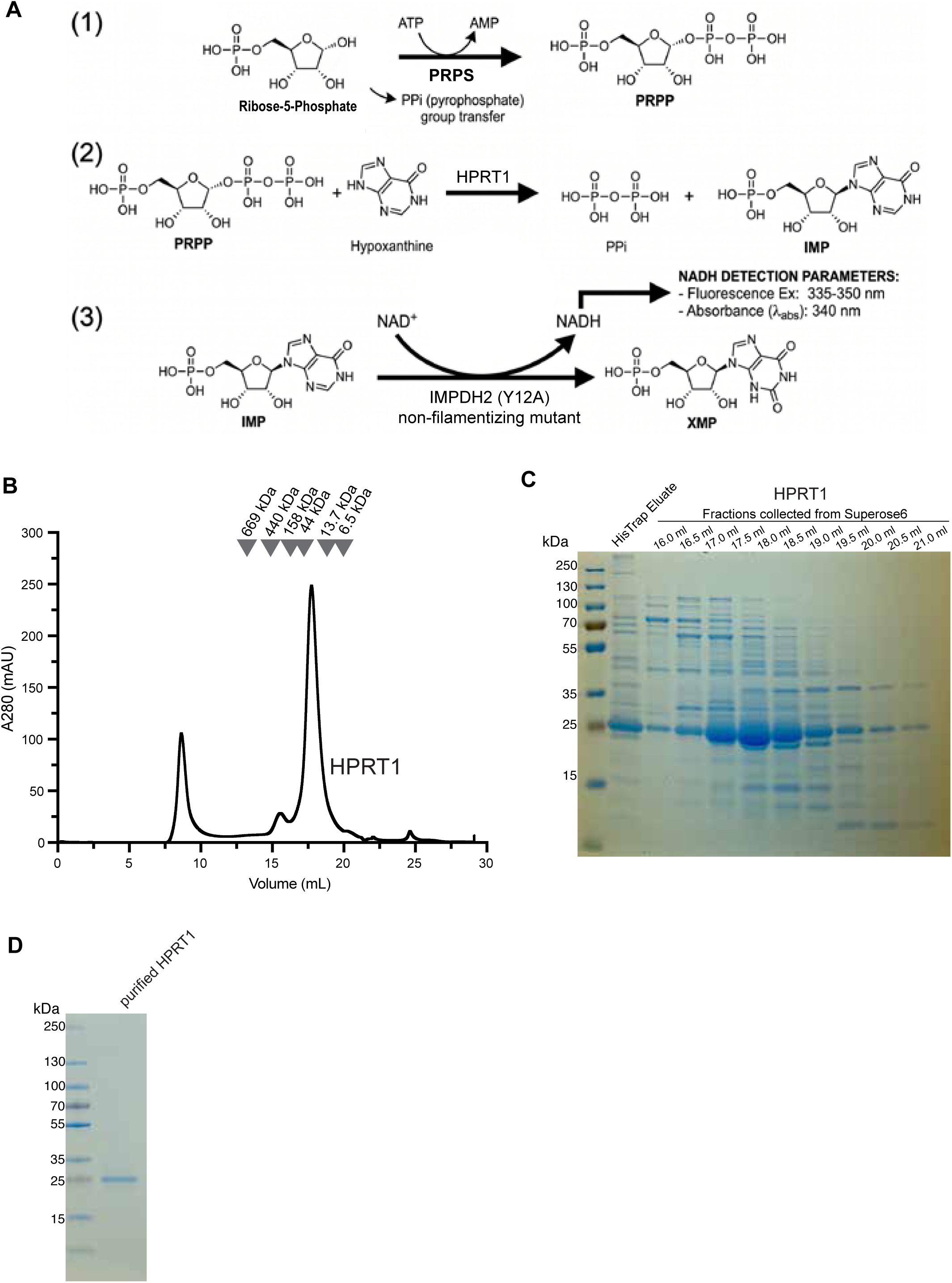
PRPS enzyme activity assay and purification of HPRT1. **(A)** Schematic of the PRPS activity assay. PRPS activity is measured through a coupled reaction in which PRPP generated by PRPS from ATP and ribose-5-phosphate is then converted by HPRT1 to IMP, followed by oxidation by IMPDH2 (Y12A; non-filamentizing mutant)^91^. This reaction produces NADH from NAD^+^, which is monitored continuously by fluorescence (excitation 335-350 nm, emission 470 nm). **(B)** Size-exclusion chromatography (SEC) elution profile of purified human HPRT1 on a Superose 6 Increase 10/300 GL column. **(C)** SDS-PAGE followed by Coomassie staining of fractions collected from the SEC run in (B), showing enrichment of HPRT1 in peak fractions. **(D)** SDS-PAGE followed by Coomassie staining of purified HPRT1 peak fraction used in activity assays.

**Supplementary Figure 20.**
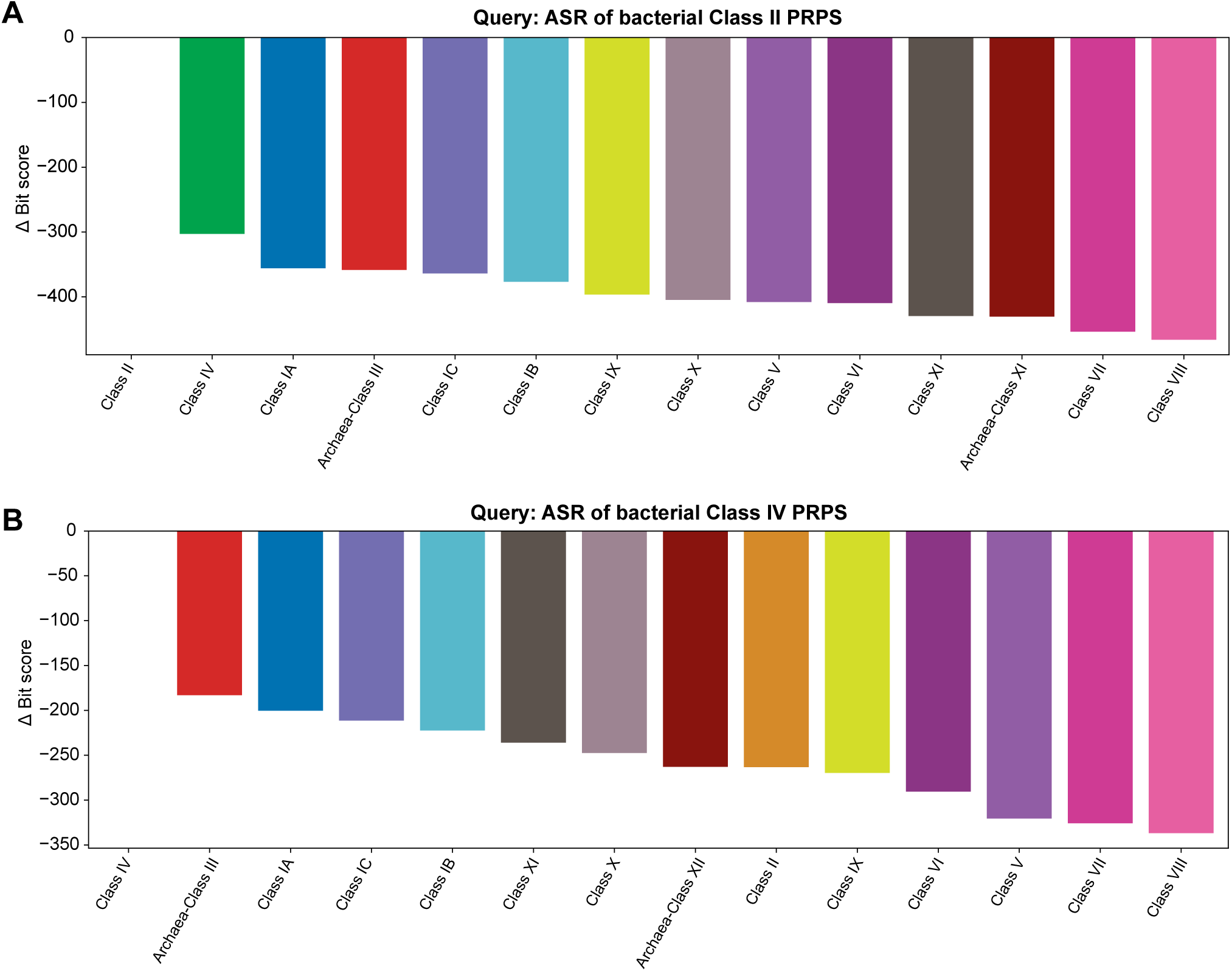
Ancestrally reconstructed bacterial sequences distinguish Class II and Class IV from other PRPS classes via HMM profiling. **(A, B)** Waterfall plots showing Δ bit scores for ancestrally reconstructed bacterial Class II (A) and Class IV (B) PRPS sequences queried against a panel of prokaryotic class-level PRPS profile HMMs. For each query, Δ bit scores are calculated relative to the top-scoring model (Δ = 0). Ancestral sequence reconstruction is described in Methods. The bacterial Class II ancestor scored highest against the Class II profile, with Class IV as the next closest class-level profile, supporting a close relationship between Class II and Class IV. In contrast, the bacterial Class IV ancestor scored highest against the Class IV profile and showed closer similarity to Class IA/Class III-associated profiles than with the Class II profile. Together, these patterns are consistent with a model in which Class IV represents an intermediate PRPS class that diverged from a Class I/III-like ancestor before subsequent emergence of Class II from a Class IV-like precursor.

