## Supplementary figures and images for "Divergent evolution of the PRPS enzymes across the tree of life"

### Supplementary File 1

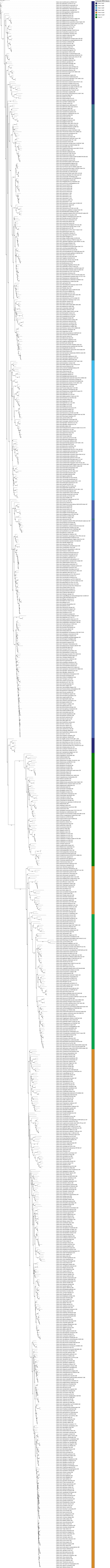
